# Environmental enrichment enhances patterning and remodeling of synaptic nanoarchitecture revealed by STED nanoscopy

**DOI:** 10.1101/2020.10.23.352195

**Authors:** Waja Wegner, Heinz Steffens, Carola Gregor, Fred Wolf, Katrin I. Willig

## Abstract

Synaptic plasticity underlies long-lasting structural and functional changes to brain circuitry and its experience-dependent remodeling can be fundamentally enhanced by environmental enrichment. It is unknown, however, whether and how environmental enrichment alters the morphology and dynamics of individual synapses. Here, we present a virtually crosstalk free, two-color *in vivo* STED microscope to simultaneously superresolve the dynamics of endogenous PSD95 of the post-synaptic density and spine geometry. With environmental enrichment, the size distributions of PSD95 and spine head sizes were sharper than in controls, indicating that synaptic strength is set more precisely with environmental enrichment. Spine head geometry and PSD95 assemblies were highly dynamic but their changes correlated only mildly. With environmental enrichment, the topography of the PSD95 nanoorganization was more dynamic; changes in size were smaller than in mice housed in standard cages and depended linearly on their original size. Thus, two-color *in vivo* time-lapse imaging of synaptic nanoorganization uncovers a unique synaptic nanoplasticity associated with the enhanced learning capabilities under environmental enrichment.

## INTRODUCTION

Over an entire lifespan, cognitive, sensory and motor learning is associated with specific changes to the cortical circuitry. Such experience-dependent plasticity is based on synaptic plasticity, plays a key role for normal brain function and apparently is neuroprotective e.g. against neurodegenerative diseases (Mandolesi et al., 2017; Nithianantharajah and Hannan, 2006). Besides such experience-dependent or extrinsic, structural changes, there is substantial evidence that synaptic structures are highly volatile intrinsically as such that synaptic connections undergo continuous spontaneous remodeling without any activity (Mongillo et al., 2017; Ziv and Brenner, 2018). Thus, intact synapses are formed despite abolishing presynaptic release (Sigler et al., 2017), or network silencing (Hazan and Ziv, 2020). Decades of neuroscience research have shown a tight correlation between structural plasticity, the morphological transformation of spines and synapses, and modifications in synaptic transmission, termed functional plasticity (Yuste and Bonhoeffer, 2001) which was confirmed recently (Holler et al., 2021). However, recent evidence suggests that not all synaptic structures follow the same dynamic and therefore structural and functional plasticity may be temporally decoupled to some extent. For example, the post-synaptic density (PSD) scaffolding proteins Homer, Shank and PSD95 increase with a delay of ~1 hour after long-term potentiation (LTP) whereas the spine heads increase in less than a minute (Bosch et al., 2014; Meyer et al., 2014). Thus, the relation between spine size and PSD organization (Arellano et al., 2007; Cane et al., 2014; Harris et al., 1992; Meyer et al., 2014) may be temporally dynamic and complex during synaptic change and the remodeling of PSD and spine head plasticity might be more independent than previously thought. Structural changes of the PSD are linked to functional changes by anchoring amino-3-hydroxy-5-methyl-4-isoxazolepropionic acid receptors (AMPAR) via transmembrane AMPAR-regulatory proteins (TARPs) to PSD95 (Compans et al., 2016; Herring and Nicoll, 2016). Both, anchoring of AMPAR and spine head remodeling by actin polymerization, are dependent on Ca^2+^ influx to the postsynapse, but rely on different intracellular signaling pathways (Chazeau and Giannone, 2016). Besides an increase or decrease in the amount of scaffolding proteins and receptors upon activity, the synapse undergoes pronounced structural changes on the time scale of minutes to hours (Cane et al., 2014; MacGillavry et al., 2013; Wegner et al., 2018; Ziv and Fisher-Lavie, 2014). Computational studies show that changes of the shape of the PSD (Franks et al., 2003) or modest clustering of AMPAR (Savtchenko and Rusakov, 2014) is a highly efficient way to modify synaptic strength without altering the total amount of receptors. Thus, a structural rearrangement on the nanoscale might be a more rapid way to induce changes in synaptic strength than incorporation of new molecules. Recently, it was shown that such a sub-synaptic nanoorganization exists in the pre- and postsynaptic side and that they are aligned in so called nano-columns (Hruska et al., 2018; Tang et al., 2016). While nanostructures of the PSD such as perforations are well observed by electron microscopy (Stewart et al., 2005), studies of the synaptic nanoplasticity (Hruska et al., 2018; MacGillavry et al., 2013; Wegner et al., 2018) and pre- and postsynaptic alignment (Hruska et al., 2018; Tang et al., 2016) rely on superresolution microscopy techniques since they are not detectable with two-photon microscopy due to the diffraction limited resolution of 300–500 nm.

Here, we use nanoscale stimulated emission depletion (STED) microscopy to map temporal changes of the spine head and PSD95 simultaneously *in vivo,* and address how changes of the PSD95 size correlate over time with spine head fluctuations at baseline that is without a dedicated stimulation paradigm. Secondly, we study whether and how activity changes the synaptic nanoorganization and plasticity. For this purpose, we employ environmental enrichment (EE) by which an animal is provided with multi-sensory stimulation, cognitive activity, social interactions and physical exercise, which modifies the degree of plasticity and dynamics of cortical sensory circuits. EE has been shown to accelerate the maturation of new neurons, enhance the expression of signaling molecules, and facilitate synaptic plasticity (Nithianantharajah and Hannan, 2006). Structural changes include increased dendritic branching and spine density in specific cortical regions (Gelfo et al., 2009; Leggio et al., 2005) and an increase in pre- and postsynaptic protein levels in major cortical brain regions (Nithianantharajah et al., 2004). However, it is unknown whether the effects of EE leave their mark only at the level of neuronal circuits and representations, while the dynamics of individual synapses remain essentially unchanged, or whether EE directly affects the dynamics and nanostructure of individual synapses, spines, and PSDs.

To unambiguously assess these relationships between spine morphology and PSD95, we set up a virtually crosstalk free two-color STED microscope for *in vivo* superresolution of EGFP and Citrine. We utilized a transcriptionally regulated antibody-like protein to visualize endogenous PSD95 without introducing overexpression artefacts (Gross et al., 2013). Together with a membrane label, we simultaneously recorded temporal morphological changes of the spine head and PSD95 nanoorganization and compared its dynamics between mice raised in EE and in standard cages (Ctr). Investigating the synaptic dynamics in the mouse visual cortex, a brain region known to exhibit enhanced plasticity under EE conditions (Baroncelli et al., 2010; Kalogeraki et al., 2017) and accessible to light microscopy, we demonstrate that spine head and PSD95 size distributions are sharpened while spine heads increase in size under EE. In addition, PSD95 underwent much stronger directional changes in size and reorganization of their nanoorganization under EE, while the average change in amplitude was smaller compared to controls. Dynamical changes of PSD95 and spine head size were only mildly correlated.

## RESULTS

### Virtually crosstalk free two-color STED microscope

To unambiguously dissect the dynamics of distinct synaptic components and examine their interactions, crosstalk free nanoscale imaging is of critical importance. We thus established a virtually crosstalk free two-color STED microscope for imaging of EGFP and EYFP or Citrine, respectively. Previous attempts featuring STED microscopy of EGFP and EYFP by two-color detection were suffering of high crosstalk requiring channel unmixing (Tønnesen et al., 2011). We therefore extended our previously described *in vivo* STED microscope (Willig et al., 2014) by an additional two-color excitation and detection to unambiguously excite the green or yellow fluorescent protein (Fig. 1A). By this means we utilize 483 nm pulses to excite EGFP and detect its spontaneous fluorescence at 498–510 nm (Det1, Fig. 1B). EYFP (or Citrine) was excited at 520 nm and detected mainly at 532–555 nm (Det2, Fig. 1B). The 483 and 520 nm excitations were switched on and off alternately while recording two consecutive line scans to temporally separate the EGFP and EYFP/Citrine emission. To determine the crosstalk between both channels, we investigated one-color labelled cells expressing EGFP or EYFP. Live-cell imaging showed that we have designed a virtually crosstalk free microscope with only ~8% crosstalk of EYFP in the EGFP channel (Channel 1) and ~5% of the EGFP signal in the EYFP channel (Channel 2) (Fig. 1C). Stimulated emission depletion is performed by a 595 nm laser beam passing a vortex phase plate to create a donut-shaped intensity profile in the focal plane of the objective as previously described (Eggeling et al., 2015).

**Figure 1:**
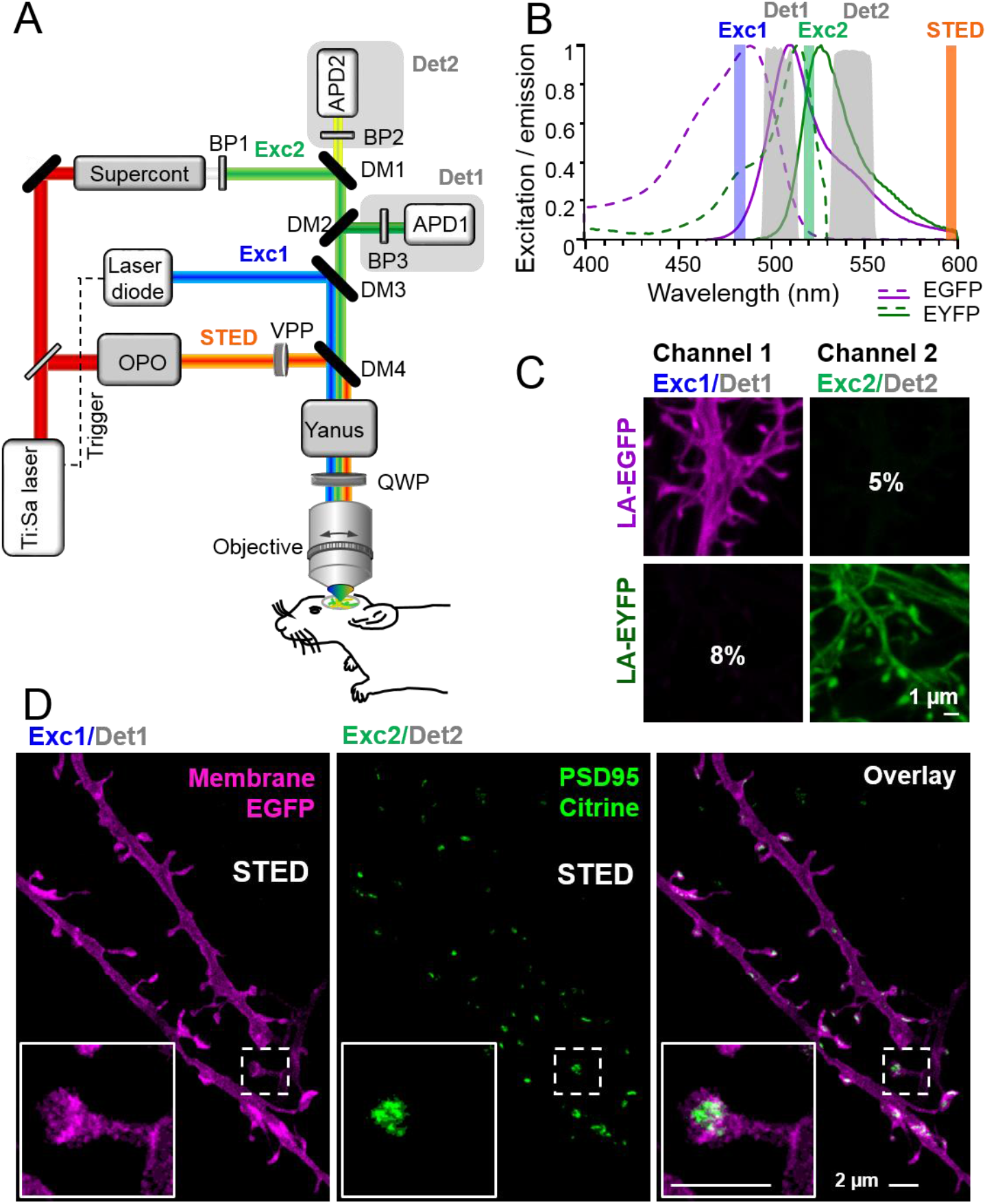
Virtually crosstalk free two-color *in vivo* STED microscopy. **(A)** Custom-designed *in vivo* STED microscope with pulsed 483 nm (Exc1) and 520 nm (Exc2) excitation and 595 nm STED laser. APD: Avalanche photon detector, BP: bandpass filter, Det: detection, DM: dichroic mirror, OPO: optical parametric oscillator, QWP: quarter wave plate, VPP: vortex phase plate. Confer Material and Methods section for details. **(B)** Excitation (dashed line) and emission (solid line) spectrum of EGFP and EYFP and wavelength regions for selective excitation and detection. **(C)** Cultured hippocampal neurons expressing the actin marker Lifeact (LA)-EGFP or LA-EYFP. Excitation of EGFP at 483 nm close to its excitation maximum (B) and detection at 498–510 nm (Det1) reduces the crosstalk to 5% in Channel 2 (C). EYFP or Citrine is excited at 520 nm close to its maximum (B) and detected at 532–555 nm (Det2) which results in a low crosstalk of 8% in Channel 1 (C). **(D)** *In vivo* STED microscopy image of apical dendrite in layer 1 of the visual cortex of an anaesthetized mouse. Labelling of membrane (myr-EGFP-LDLR(Ct)) and PSD95 (PSD95-FingR-Citrine) visualizes the spine morphology at superresolution and PSD95 nanoorganization (D, inset). Images are smoothed and represent maximum intensity projections (MIPs). No unmixing is performed due to the low crosstalk.

### Two-color in vivo STED microscopy of endogenous PSD95 and spine morphology

Having established the virtually crosstalk free detection of two STED approved fluorescent proteins (Nagerl et al., 2008; Willig et al., 2006), we set out to simultaneously superresolve the nanoorganization of the post-synaptic density protein PSD95 and the associated spine morphology *in vivo*. To this end, we generated recombinant adeno-associated viral (AAV) particles encoding fusion proteins under control of the human Synapsin promoter (hSyn). To visualize PSD95, we expressed the transcriptionally regulated antibody-like protein PSD95-FingR, that has been shown to label endogenous PSD95 (Gross et al., 2013; Willig et al., 2021) attached to the fluorescent protein Citrine. For spine morphology, we expressed myr-EGFP-LDLR(Ct), a combination of a myristoylation site (myr), EGFP and the C-terminal (Ct) cytoplasmic domain of low density lipoprotein receptor (LDLR), a potent marker for the dendritic membrane (Kameda et al., 2008; Willig et al., 2021). In order to reduce the density of labelled neurons, the fluorescent labels were incorporated into AAVs in reverse orientation in the vector between double-floxed inverted open reading frames (DIO). Expression of the labels was enabled by Cre recombinase encoding AAVs co-injected at low concentration (Willig et al., 2021). Hence, the density of labeled neurons was tuned by the dilution of the Cre expressing AAV and the brightness of the PSD95 and membrane labelling by the concentration of the respective AAVs; thus the density of labelled cells and their brightness were independent variables. We co-injected the three AAVs into pyramidal cell layer 5 of the visual cortex. Three to six weeks after transduction, the mice were anaesthetized and a cranial window was inserted over the visual cortex as described before (Steffens et al., 2020). STED microscopy visualized dendrites, spines, and attached PSD95 assemblies in layer 1 of the visual cortex at superresolution (Fig. 1C) with a resolution of at least ~70 nm for Citrine and ~84 nm for EGFP (Fig. S1A, B). The inset in Fig. 1D reveals a perforated nanoorganization of endogenous PSD95 that would not be detectable with conventional *in vivo* light microscopy (Fig. 1D) due to the insufficient resolution. Due to the low crosstalk, a computational post-processing such as unmixing was not required. We observed mainly dendrites that expressed both, EGFP and Citrine, and only rarely found dendrites expressing either Citrine or EGFP alone.

### EE housed mice exhibit a sharper size distribution of spine head and PSD95 assemblies while their heads are larger than for Ctr

Having established the two-color *in vivo* STED microscope, we addressed the question of whether spine or post-synaptic density exhibit systematic differences in size between EE and Ctr mice. EE mice were housed in a large, two-floor cage equipped with three running wheels for physical exercise, a labyrinth for cognitive stimulation, a tube and a ladder to change between the two levels, and were kept in groups of up to 12 female mice to allow manifold social interactions (Fig. S2). Control mice were raised in pairs in standard cages without any equipment (Fig. S2). The mice were transduced with AAVs as described above and 3–6 weeks after transduction, a cranial window was implanted above the visual cortex. We acquired two-color z-stacks mainly of dendrites, which were in parallel to the focal plane. To analyze differences in morphology of the dendritic spines, we encircled the spine head and PSDD95 at their largest extent and computed the respective area (Fig. 2A, cf. section Methods for details). We only considered spines containing a PSD95 label and therefore most likely form functional synapses. Occasionally we observed long, thin spines without a head, sometimes called filopodia in the literature which were not analyzed. The spine head area correlated with the PSD95 area for EE and Ctr (Fig. 2B), corroborating EM and two-photon microscopy studies (Arellano et al., 2007; Cane et al., 2014; Meyer et al., 2014). The histograms of spine head (Fig. 2C) and PSD95 area (Fig. 2D) were positively skewed and the distributions were significantly different between EE and Ctr mice. We determined a median spine head area for EE housed mice of 0.527 (interquartile range (IR): 0.359–0.844) μm^2^ and 0.462 (IR: 0.290–0.787) μm^2^ for Ctr mice. The area of PSD95 on the same set of spine heads was 0.158 (IR: 0.107– 0.269) μm^2^ in median for EE mice and 0.161 (IR: 0.092–0.286) μm^2^ for Ctr mice which is in accordance with previous EM studies in the mouse visual cortex (Harris and Weinberg, 2012). To further dissect these differences, we next plotted the histograms of the logarithm of spine head (Fig. 2E) and PSD95 areas (Fig. 2F). All four histograms (Fig. 2E, F) are symmetric and well described by a Gaussian function. This is in line with prior work showing that the spine and PSD95 fluorescence is log-normally distributed in general which is predicted by multiplicative processes of ongoing spine plasticity (Hazan and Ziv, 2020; Loewenstein et al., 2011). The Gaussian function fitting the spine head area is shifted to the right and is narrower for EE housed mice (Fig. 2E). This is manifested by a significantly larger spine head area (Fig. 2G) and smaller variance of the size distribution (Fig. 2I) of EE housed mice. The distributions of the logarithm of the PSD95 area (Fig. 2F) are almost centered and thus the average area is not significantly different between EE and Ctr mice (Fig. 2H). However, the variance (Fig. 2J) is smaller for EE housed mice which is also visible in the Gaussian distribution which is narrower for EE housed mice (Fig. 2F). Since previous studies often used the brightness of PSD95 assemblies or spines, respectively, as a measure of size, we also analyzed the PSD95 brightness and found a coefficient of determination R^2^ = 0.76 between the brightness and nanoscale size (Fig. S3A). Spine density was not significantly different between EE and Ctr housed mice (Fig. S3B). These results show that the enriched environment leaves a trace in spine head size and variance of the post-synaptic size, both of which have been shown to correlate with synaptic strength (Holler et al., 2021).

**Figure 2:**
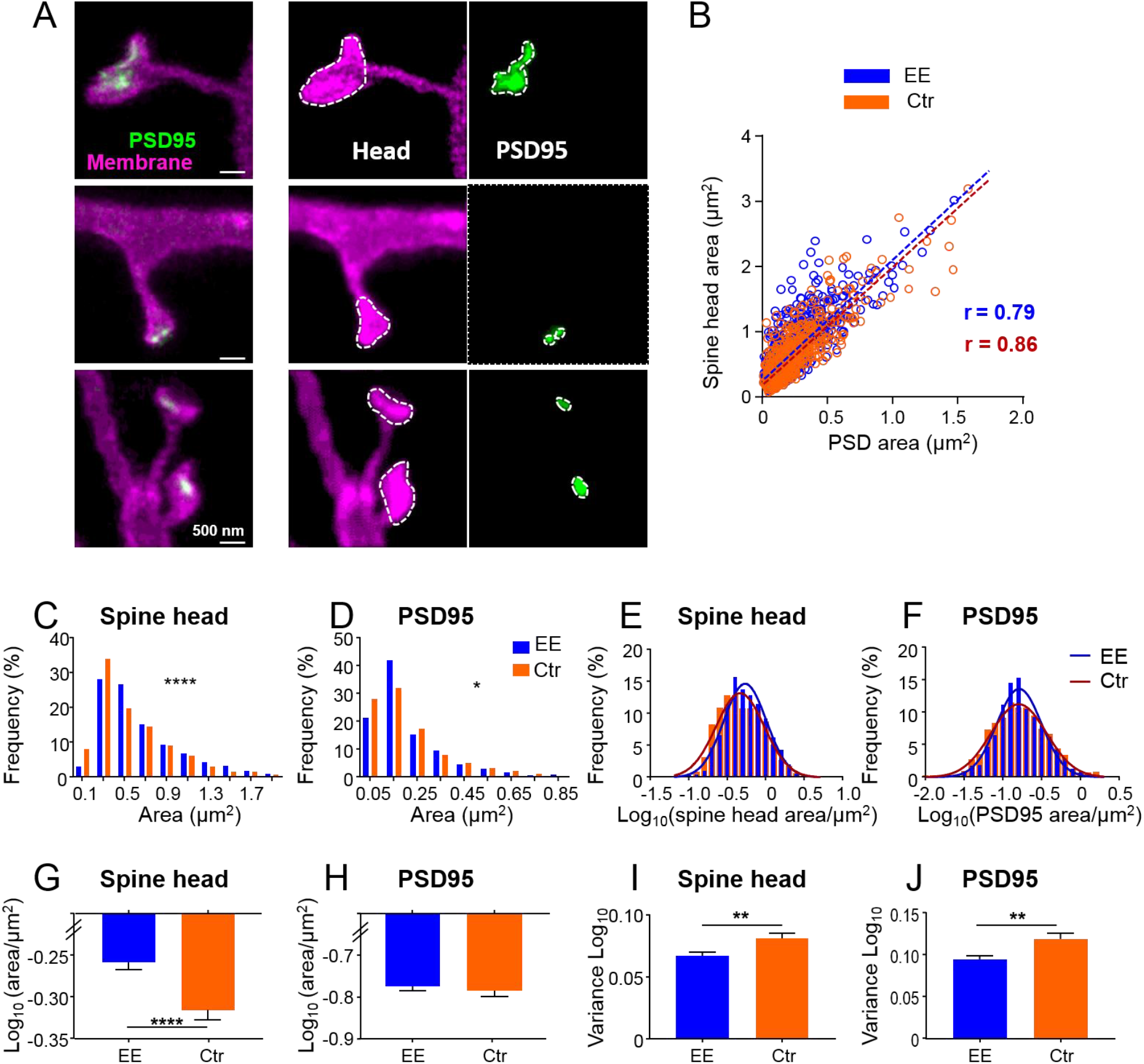
The size distributions of the spine head and PSD95 area are sharper and show larger heads for mice housed in environmentally enriched (EE) than standard (Ctr) cages. **(A)** STED images of dendritic spines (magenta) and associated PSD95 assemblies (green). Images are smoothed; MIP (left). Smoothed and contrast enhanced images for area analysis (middle, right). Spine heads (middle) and PSD95 assemblies (right) were encircled to compute the area. **(B)** Strong correlation of absolute spine head and PSD95 area in Ctr (orange) and EE (blue) housed mice. Linear regression lines are dashed and Pearson’s correlation coefficient *r* displayed (EE and Ctr, deviation from zero: p < 0.0001). **(C, D)** Frequency distributions of spine head area (C) and PSD95 area (D) are positively skewed and significantly different between EE and Ctr housed mice (Kolmogorov-Smirnov test, C: ****p < 0.0001, D: *p = 0.013). Graphs display center of BIN, single large values are cut off. **(E, F)** Same data as shown in (C, D), but logarithmic values. Solid lines represent Gaussian functions fitted to the respective histogram. **(G–J)** Mean (G: EE_spine_: −0.259, Ctr_spine_: −0.317, H: EE_PSD_: −0.771, Ctr_PSD_: −0.781) and variance (I, J) of logarithmic data shown in (E, F) + SEM (Unpaired *t*-test with Welch’s correction: G: ****p < 0.0001, H: p = 0.54, I: **p = 0.006, J: **p = 0.003). Number of analyzed mice and spines: EE: 4x ♀-mice, n_Spine/PSD95_ = 795; Ctr: 4x ♀-mice, n_Spine/PSD95_ = 634.

### Correlated changes of PSD95 and spine head area over minutes to hours

Above we showed a strong correlation between the size of PSD95 assemblies and the spine head (Fig. 2B) which corroborates previous findings of a close structural correlation between PSD and spine size. However, whether dynamic changes between these two features are also strongly linked has remained unknown. We thus asked whether and how temporal changes of these parameters were correlated. Therefore, we performed time-lapse STED microscopy of spine morphology and PSD95 for EE and Ctr housed mice for up to 4 hours as described above and analyzed temporal changes of these parameters (Fig. 3A–K). We recorded STED images at different fields of view for three time points at varying time intervals Δt of 30 min, 60 min or 120 min (Fig. 3A, B) and analyzed the size of the spine head and PSD95 assembly as described above. The average spine head area and PSD95 area did not show large changes over the time period of 120 min (Fig. 3C). Thus, we did not observe directed changes as a result of light induced stimulation or phototoxicity and neither observed blebbing. We computed normalized size changes over time so that both positive changes (growth) and negative changes (shrinkage) were symmetric with a boundary value of ±1. Figure 3D–F shows scatter plots of the normalized changes of the spine head over changes of the PSD95 area. For each time interval the data points are scattered over all 4 quadrants (for percentage changes refer to Fig. S4A–C). This means we observed positively correlated changes of growing spine heads and growing PSD95 areas (or shrinking, respectively) (Fig. 3J) as well as anti- or negatively correlated changes, i.e. shrinking PSD95 areas on a growing spine or vice versa (Fig. 3K). A linear regression analysis revealed a significant positive correlation (*r* between 0.15 and 0.39, Fig. 3D–F), such that an increase of PSD95 area is more frequently accompanied by an increase in spine head area and vice versa. Interestingly, PSD95 assemblies display larger changes in size than spine heads when normalized which is evident from the elliptic distributions in Figure 3D–F or the median percent changes (Fig. S4D, E); spine heads grow by ~20% and shrink by ~15% while PSD95 assemblies grow by ~20–25% and shrink by 25–30%. Next, we asked whether changes were stronger for EE or Ctr housed mice. Therefore, we computed the total variance for spine head and PSD95 area changes (Fig. 3G) and found a higher variance for Ctr housed mice than EE after each time interval which was significantly increased after 60 min. This total variance was rather constant over all different time intervals for both EE and Ctr housed mice. To dissect whether this difference in variance was due to correlated or anti-correlated changes, we performed a principal component analysis (PCA). Principal component 1 (PC1) is by definition the axis along which the data shows the maximum of variance and the eigenvalue of PC1 is the variance along this axis. The PC1 vector was for all three time intervals in the first (and third) quadrant (Fig. 3D–F) and therefore attributing for correlated changes. The second principal component (PC2) is by definition perpendicular to PC1 and therefore in the second (and fourth) quadrant (Fig. 3D–F). The variance of both components, PC1 (Fig. 3H) and PC2 (Fig. 3I), was higher for Ctr than EE house mice after each time interval; this increase was statistically significant for the 60 min interval for PC1 and for the 30 and 60 min intervals for PC2. The variance of PC1 is ~3 times larger than that of PC2 (Fig. S4F).

**Figure 3:**
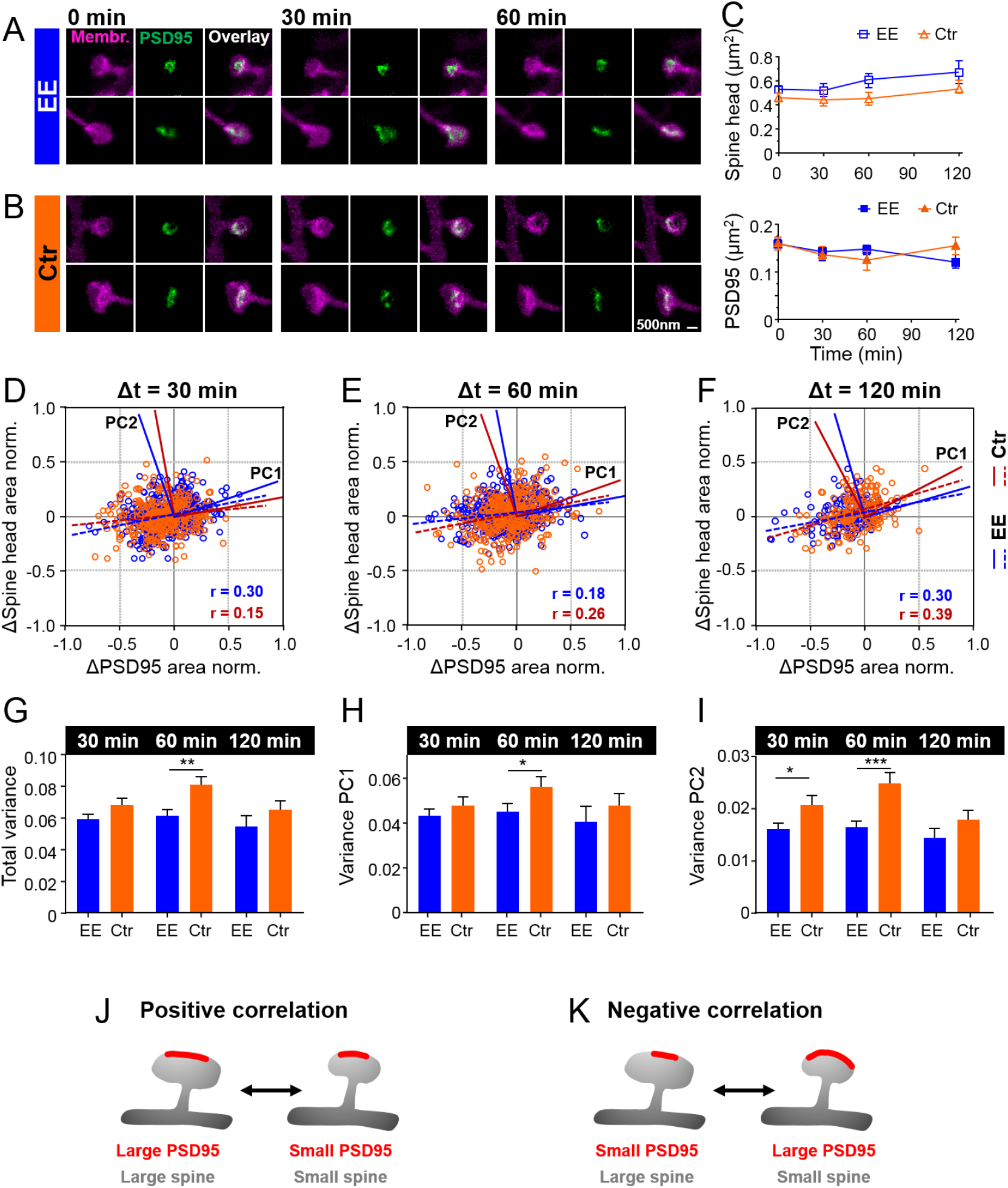
Temporal changes of spine head and PSD95 area are positively and negatively correlated and larger for Ctr than EE house mice. **(A, B)** Representative sections of spine heads and corresponding PSD95 of time-lapse *in vivo* two-color STED microscopy images for EE (A) and Ctr (B) housed mice at time points of 0 min, 30 min and 60 min. Images are smoothed and shown as MIP. **(C)** Median ±95% confidence interval (CI) of spine head and PSD95 areas of Ctr (triangle, orange) and EE (square, blue) housed mice over time. **(D–F)** Normalized changes in spine head and PSD95 area after 30 min, 60 min and 120 min time intervals. Linear regression lines are dashed (deviation from zero: p < 0.0001) and Pearson’s correlation coefficient *r* is displayed. Solid lines represent the principal components 1 and 2 (PC1, PC2) of the principal component analysis (PCA) for Ctr (red) and EE (blue), respectively. **(G–I)** Variance of normalized changes of PSD95 assemblies and spine head area plotted in (D–F); mean of total variance (G), variance of PC1 (H) and PC2 (I) +SEM (Unpaired t-test EE vs. Ctr; G: 30 min: p = 0.088, 60 min: **p = 0.0015, 120 min: p = 0.23; H: 30 min: p = 0.36, 60 min: *p = 0.045, 120 min: p = 0.40; I: 30 min: *p = 0.027, 60 min: ***p < 0.001, 120 min: p = 0.17). **(J, K)** Illustration of temporal changes between spine head size and PSD95 area; Growth and shrinkage of spine head size and the area of PSD95 assemblies goes hand-in-hand, i.e. are positively correlated (J). An anti- or negative correlation is observed for a growing PSD95 assembly on a shrinking spine and vice versa (K). (C–I) Number of analyzed mice and spines: 4x ♀-mice for EE and 4x ♀-mice for Ctr; number of spines with PSD95 assemblies: EE, t = 0: 795, t = 30 min: 326, t = 60 min: 388, t = 120 min: 133; Ctr, t = 0: 634, t = 30 min:233, t = 60 min: 285, t = 120 min: 189.

In summary, the synaptic structure is highly dynamic and spine heads and PSD95 assemblies change in size by > 20% already after 30 min. The amplitude of changes is similar for all time intervals up to 120 min and is higher for Ctr than EE housed mice. These results suggest that Ctr housed mice undergo stronger morphological changes.

### Multiplicative downscaling of PSD95 area is different between EE and Ctr housed mice

Since our parameters were log-normally distributed, indicating a multiplicative dynamic, we asked whether the temporal changes were also regulated by multiplicative processes (Hazan and Ziv, 2020; Loewenstein et al., 2011). Thus, we plotted the synaptic size after different time points as a function of the original size (Fig. 4A–F). A linear regression analysis revealed a slope <1 and positive y-intercept for spine head and PSD95 area, for both EE and Ctr housed mice. Plotting the size difference between the different time points as a function of the original size indicates that large spines tend to shrink and small spines tend to grow; this becomes evident as the changes are rather negative for larger sizes and positive for smaller sizes (Fig. S5). A recent study uses a Kesten process as a model for spine dynamics and their skewed size distribution. In this model a noisy multiplicative downscaling is combined with a noisy additive term (Hazan and Ziv, 2020; Statman et al., 2014). Thus, the negative slope as shown in Fig. S5 which is equal to 1-slope of Fig. 4 would be regarded as a time-dependent multiplicative downscaling factor. We observed a significantly different slope, i.e. multiplicative downscaling for the PSD95 area between EE and Ctr housed mice (Fig. 4F). While the slope was constant or decreased slightly over time for EE housed mice it raised up to nearly 1 for Ctr mice. Interestingly, this effect was specific for PSD95 and was not observed for the spine head size (Fig. 4A–C). The slope for spine head size changes was rather constant over time and similar between EE and Ctr mice. In summary, we found a significant increase in downscaling over time for PSD95 in EE mice compared to Ctr after 60 min. Comparing a silenced network with an active network *in vitro* Hazan *et al.* found a less pronounced downscaling of synaptic sizes over time after silencing (Hazan and Ziv, 2020). Although these experiments were performed *in vitro*, the more pronounced downscaling, i.e. smaller slope, we find for EE housed mice shows the same tendency and might be explained by a higher activity level in EE mice and also potentially cause the narrower size distribution of EE housed mice (Fig. 2F, J) (Hazan and Ziv, 2020). This pronounced dynamics might restore fluctuations away from the average parameters in EE in line with a narrower and thus sharper size distribution of PSD95 assemblies and spine heads that are also associated with EE. However, a downscaling factor of 0.7–0.8 which we found for PSD95 in EE housed mice was observed by Hazan and Ziv only after about 20 hours.

**Figure 4:**
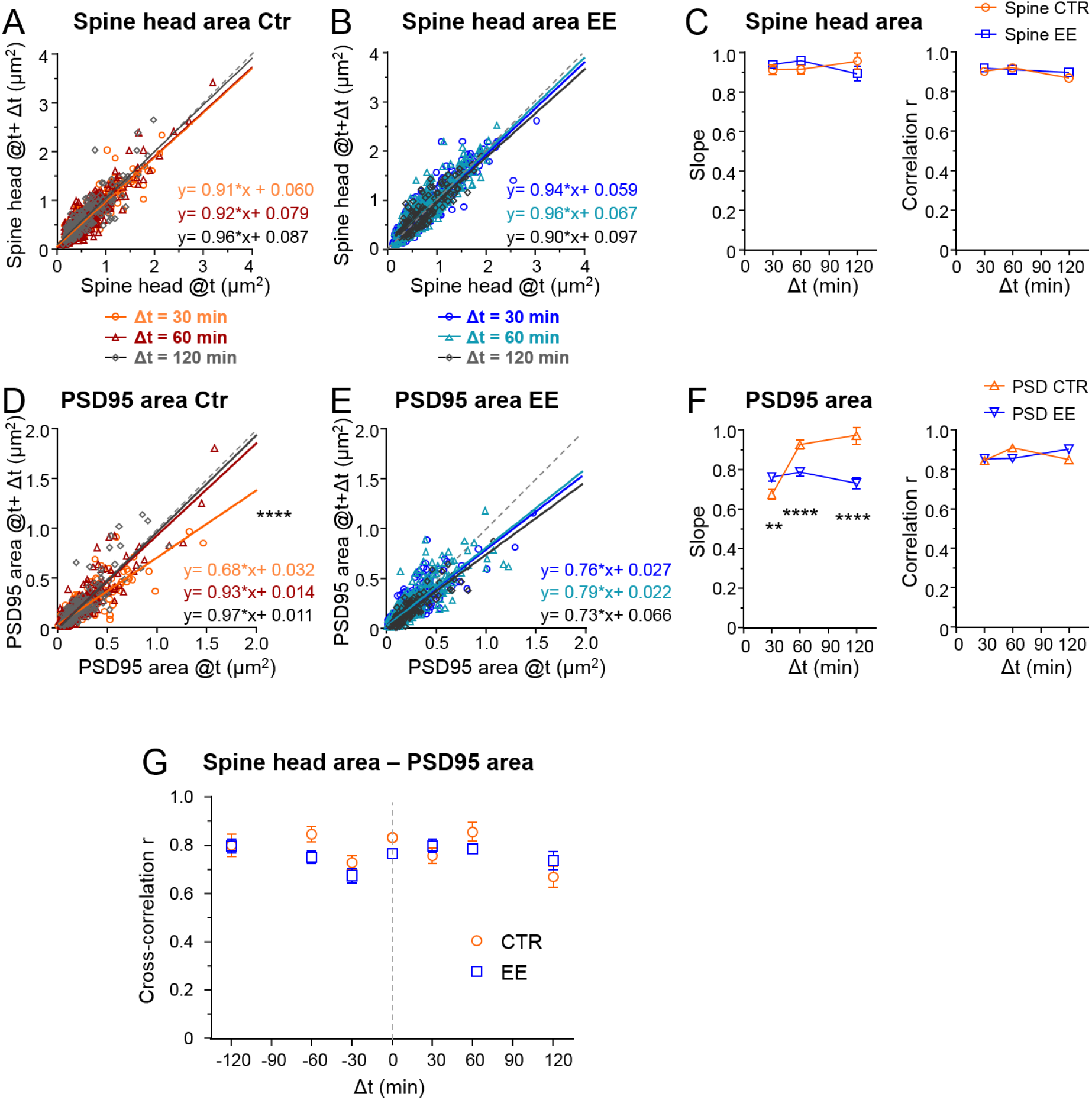
EE housed mice show an increase in multiplicative downscaling of PSD95 area over Ctr. **(A– F)** Spine head and PSD95 area after different time intervals *Δt* of 30 min, 60 min and 120 min as function of its area at time *t* (A, B, D, E). Solid lines show linear regression fits of the displayed equation; the identity line is dashed (Analysis of Covariance; are slopes equal? A: p = 0.53, B: p = 0.29, D: ****p < 0.0001, E: p = 0.39). (C) Slope ±SE of fit to spine head changes (A, B)(left, are slopes different? EE vs. Ctr: 30 min: p = 0.37, 60 min: p = 0.11, 120 min: p = 0.26) and Pearson’s correlation *r* (right) of linear regression. (F) Slope ±SE of fit to PSD95 area changes (D, E)(left, are slopes different? EE vs. Ctr: 30 min: **p < 0.01, 60 min: ****p < 0.0001, 120 min: ****p < 0.0001) and *r* value (right) (). **(G)** Cross-correlation of spine head and PSD95 area for EE and Ctr housed mice. Error bars are bootstrap ±SD.

### Despite fluctuations spine heads and PSD95 assemblies are stable in average over 2 hours

Next, we asked how these large temporal changes influence the mean size. Do growing spines continue to grow and do shrinking spines continue to shrink? Thus, we computed the Pearson’s correlation value *r* which was relatively constant over all time intervals and not different between spine heads and PSD95 assemblies or EE and Ctr house mice (Fig. 4C, F, right). As such we did not find changes in its *r* or its corresponding R^2^ value such as described *in vitro* (Hazan and Ziv, 2020) at our time window of up to 2 hours. This indicates that large spine heads tend to stay rather large and smaller ones small. To analyze the coupling between spine head and PSD95 area over time, we computed the cross-correlation between these measures. Fig. 4G shows that the cross-correlation between PSD95 and the spine head was ~0.8 over all time intervals of 0–120 min and thus we did not observe directional changes of these parameters. In particular, we did not observe that PSD95 increase would systematically precede a subsequent spine head increase or vice versa as both would result in an asymmetric cross-correlation.

### PSD95 nanoorganization changes faster in EE mice

Previous work with EM has shown that PSDs on large spines, often called mushroom spines, are frequently perforated (Stewart et al., 2005). In our previous publication (Wegner et al., 2018), we investigated PSD95 morphology in homozygous knock-in mice and showed that PSD95 assemblies on large spines were also often perforated and appeared ring-like or clustered. This nanoorganization was highly dynamic and changed in shape within a few hours. Now, we asked whether these changes would be different for mice housed in EE. Therefore, we selected perforated PSD95 assemblies and investigated their morphological changes analogously to the study by Wegner *et al.* (Wegner et al., 2018). Fig. 5A shows examples of two-color STED images of perforated PSD95 assemblies and their associated spine revealing temporal changes in the PSD95 nanoorganization. Morphological changes were categorized for each time point as no change, subtle change, or strong change with respect to the first observation at t = 0 by three independent persons (Fig. 5B, C). Ctr housed mice showed a clear trend: the longer the measurement interval, the greater the morphological changes (Fig. 5C); the percentage of PSD95 nanoorganization showing no change decreased from ~40% to ~10%, while the percentage for strong changes nearly tripled for Ctr. For EE housed mice, we observed similar, strong changes within all investigated time intervals; ~20% of the PSD95 assemblies did not change at all and ~80% underwent a change at all time points (Fig. 5B). Although the percentage changes with respect to t = 0 were similar for different time intervals, it should be noted that this does not exclude dynamic changes between the time points. Thus, changes in Ctr housed mice increased over the observation period and reached a level similar to that of EE housed mice after 120 min. This suggests that the PSD95 nanoorganization is more dynamic in EE mice than in mice raised under control conditions within the investigated time course. Now, we asked whether the different dynamic in morphological changes would be reflected as well in a different nanoorganization. Therefore, we categorized the nanoorganizations into perforated (a continuous shape with a sub-structure such as ring- or horseshoe-like) or clustered. Fewer nanoorganizations of EE housed mice were perforated and a larger fraction showed three or more clusters compared to Ctr mice (Fig. 5D). This suggests that the PSD95 nanoorganization might play a substantial role in synaptic remodeling and is specifically shaped by experience or activity.

**Figure 5:**
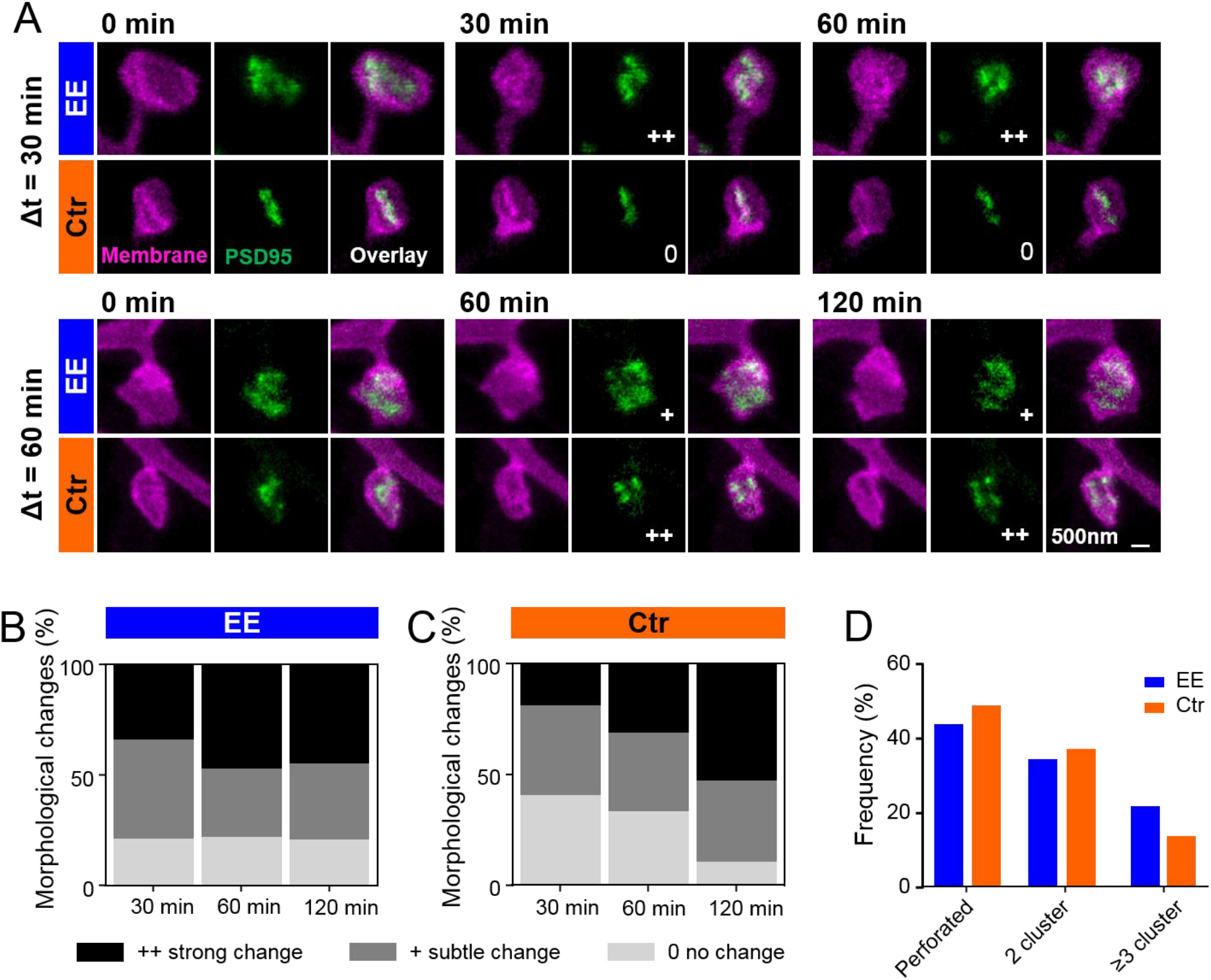
PSD95 nanoorganization is different between EE and Ctr and changes faster for EE housed mice. **(A)** Sections of two-color STED images (smoothed, MIP) of EE and Ctr housed mice at the indicated time points showing the spine membrane (magenta) and PSD95 (green). Morphological change is indicated with: **0** = no change; **+** = subtle change; **++** = strong change. **(B, C)** Stacked histogram of the relative frequency of morphological changes in PSD95 nanoorganization in EE (B) and Ctr (C) housed mice; all changes refer to t = 0 min. **(D)** Morphometry of PSD95 nanoorganization of EE and Ctr housed mice at t = 0. (B–D) Number of analyzed PSD95 assemblies in EE: 30 min: n = 38, 60 min: n = 68, 120 min: n = 29; Ctr: 30 min: n = 27, 60 min: n = 48, 120 min: n = 38.

## DISCUSSION

Interactions with the environment play a key role in restructuring and refining the neuronal circuitry in the brain. We here used time-lapse *in vivo* STED microscopy, a superresolution light microscopy technique with nanoscale resolution, to study experience-dependent changes of the synaptic nanoorganization as well as its nanoplasticity in the intact brain. We designed a two-color, virtually crosstalk free *in vivo* STED microscope with a resolution of 70–80 nm for simultaneous imaging of spine morphology and PSD95 in the visual cortex of anesthetized mice. We found a significantly smaller variance in size of the spine head and PSD95 nanoorganization in EE mice while the average spine head size was increased compared to Ctr mice. Both parameters were highly volatile and we observed an average growth of >20% and shrinkage of >18% for spine heads and PSD95 assemblies within 30 min while changes in PSD95 assembly size were slightly larger than those of the spine head. About 3/4 of these changes were positively correlated and ~1/4 showed a negative correlation; Ctr mice exhibited a larger variance of changes than EE housed mice. All parameters exhibited multiplicative downscaling which was significantly different for PSD95 assembly size between EE and Ctr mice. Dynamical rearrangement of the nanostructure was faster in EE housed mice.

### Two-color STED nanoscopy of synaptic nanoorganization

Previous studies mainly used two-photon microscopy to study synapse and spine plasticity *in vivo* and *in vitro* (Bosch et al., 2014; Cane et al., 2014; Meyer et al., 2014; Villa et al., 2016). Due to its limited optical resolution of 300–500 nm, spine brightness is often taken as a measure of size (Cane et al., 2014; Hofer et al., 2009; Matsuzaki et al., 2004). This is not very accurate since the brightness can be influenced by many factors such as expression level of the fluorescence label, the excitation laser intensity, group velocity dispersion or scattering in the tissue. Therefore, spine brightness is typically normalized to the brightness of the dendrite, making comparative studies and determination of absolute sizes difficult. To circumvent the diffraction limit two-photon microscopy was combined with STED to achieve nanoscale resolution (Moneron and Hell, 2009; Panatier et al., 2014). However, a drawback of this approach is the relatively high crosstalk when using two colors. Thus, the most commonly used EGFP and EYFP have a crosstalk of as much as 92% and 27% in the respective other channel, requiring linear unmixing (Tønnesen et al., 2011). Unmixing works well for structures which are relatively bright and spatially separated such as for volume labeled spines and astrocytes (Panatier et al., 2014) but can produce artefacts in dark and overlapping structures such as the post-synapse and spine we were studying in this project. Therefore we used an approach from two-color STED with organic dyes in the red emission spectrum (Bottanelli et al., 2016; Göttfert et al., 2013) and designing a two-color STED microscope with two different excitation lasers (one-photon excitation) and two spectrally separated detection channels. This approach gave us an almost negligible crosstalk of 5% for EGFP and 8% for Citrine rendering unmixing obsolete.

### Temporal changes of PSD95 assemblies and spine heads are largely uncorrelated

We used the transcriptionally regulated antibody-like PSD95-FingR, which efficiently labels endogenous PSD95 (Cook et al., 2019; Gross et al., 2013) and excludes artefacts which appear after PSD95 overexpression (El-Husseini et al., 2000; Stein et al., 2003). While we observed a substantial correlation between the absolute spine head and PSD95 assembly size (*r* ~ 0.8, Fig. 2B), corroborating previous EM studies (Arellano et al., 2007; Harris et al., 1992), we found only weak correlations between their temporal changes (*r* ~ 0.2–0.4, Fig. 3D–F). Most of the changes were positively correlated while about ~1/4 was attributed to negatively correlated changes. LTP experiments have shown a rapid increase in spine head size and AMPA receptor current after potentiation but a delayed increase of PSD95 which was not increased in amount until a late protein synthesis-dependent phase of LTP after ~60 min (Bosch et al., 2014; Meyer et al., 2014). Although these experiments were performed in cell culture and with PSD95 overexpression, they indicate that spine and PSD enlargement might not be temporally concerted and therefore regulated differently (Compans et al., 2016; Herring and Nicoll, 2016). Thus, the largely uncorrelated changes we observed at baseline *in vivo* at similar time scales of 30–120 min support a temporal disentanglement of synapse and spine morphology. On the other hand we found no directional changes that would demonstrate that the PSD95 size systematically follows an increase in spine head size since the cross-correlation is largely symmetric (Fig. 4G).

But how can the time-line of AMPA receptor incorporation and PSD95 increase be so different although AMPA receptors are linked to PSD95 by TARPs? A slot model proposes that the actin polymerization during LTP increases the number of slots which can harbor AMPA receptors (Herring and Nicoll, 2016). Thus, the increase of AMPA receptors would not depend on an increase of the amount of PSD95 proteins at the synapse but on an activation of binding sites of AMPA receptor to the PSD95 by a still unknown mechanism.

The changes in spine head are very similar in amplitude to the changes we have reported recently for long-term changes in the cortex of a transgenic mouse (Steffens et al., 2021). The maximum change we observed for spine head area changes was a growth of ~ 200%. This is in a similar range as reported for spine or spine head volume increases after glutamate uncaging (Bosch et al., 2014; Meyer et al., 2014; Tønnesen et al., 2014) and might reflect potentiation of single spines. Changes in the amount of PSD95 after glutamate uncaging, however, were reported always at much lower fractions than the spine volumes (Bosch et al., 2014; Meyer et al., 2014) and thus it is interesting that we observe a slightly larger increase of PSD95 assembly area than spine head area *in vivo*. However, a detailed assessment of the synaptic nanophysiology is still missing.

### Synapses and spines of EE housed mice are sharper defined in size than Ctr

To study activity-dependent changes, we compared adult mice housed in standard cages with age matched EE housed mice and analyzed synaptic morphology and plasticity in the visual cortex after the critical period. We found a significantly smaller variance and thus narrower distribution of spine head and PSD95 area in EE housed mice. At the moment different models of how neuronal networks adapt to changes in activity to undergo homeostatic plasticity are considered (Lee and Kirkwood, 2019). In the model of synaptic scaling, an increase in activity leads to a downscaling of excitatory synapses. The downscaling is multiplicative and therefore changes affect the average as well as broadening of the size distribution as shown, for example, in a silenced network (Hazan and Ziv, 2020). The narrower size distributions in EE housed mice which experience enhanced activity are therefore in line with such a model and a demonstration of synaptic adaptation *in vivo*. However, we observed an increase in spine head size after enrichment and no significant difference between PSD95 assembly size; therefore the changes are not pure synaptic scaling. This is in line with recent *in vivo* observations that in the adult cortex homeostatic plasticity is rather input-specific and not multiplicative for the whole ensemble of synapses (Lee and Kirkwood, 2019).

### Stronger multiplicative downscaling in EE housed mice for PSD95

Changes of spine heads and PSD95 assemblies correlated with their absolute size for EE and Ctr house mice. Small spines tended to grow and large spines tended to shrink. Such morphological changes can be modeled by a Kesten process including additive and multiplicative changes. This model entails a time-dependent multiplicative downscaling factor (Hazan and Ziv, 2020) corresponding to the slope in Fig. 4A–F. Silencing a neuronal network not only resulted in an average size increase and broader distribution but also in a weaker multiplicative downscaling (Hazan and Ziv, 2020). Our observations are in line with these results in such that the increase in activity by enrichment strengthened the multiplicative downscaling over time for PSD95 after 60 and 120 min (Fig. 4F). This in turn might explain the narrower size distribution according to the model by Hazan and Ziv (Hazan and Ziv, 2020). However, we did not observe significant changes for multiplicative downscaling for the spine head, neither between EE and Ctr nor between different time intervals. We note in passing that it should not be surprising that our *in vivo* experiments do not completely match expectations from artificially silenced neuronal culture experiment. As discussed above, there is evidence that homeostatic plasticity is not global but input-specific (Lee and Kirkwood, 2019).

### Enhanced nanoplasticity by experience

With STED nanoscopy we often observed a perforated nanoorganization of PSD95 undetectable by two-photon or conventional imaging. The pattern of the PSD95 nanoorganization was different and structural changes of the nanopattern occurred more rapidly in EE housed mice, suggesting greater synaptic flexibility of EE housed than Ctr housed mice. Accumulating evidence indicates that pre- and postsynaptic structures are patterned and often clustered. The functional consequences of this nanopattern, however, are not fully understood. An emerging view is that clusters are trans-synaptically aligned in ‘nanocolumns’ which are organized by activity (Chen et al., 2018). Moreover, computational simulations predict that changes in the shape of the PSD are a way to align post-synaptic receptors to pre-synaptic release sights (Franks et al., 2003; Savtchenko and Rusakov, 2014). This implies that a reorganization of receptors might alter synaptic strength – independently of changes in the amount of receptors (Chen et al., 2018). Our finding of an increased structural plasticity of the PSD95 nanoorganization but not increased average size in EE mice might therefore reflect the reported increase of synaptic plasticity after enrichment (Artola et al., 2006; Buschler and Manahan-Vaughan, 2012).

### Outlook

Our results demonstrate that experience influences the synaptic nanopattern and facilitates structural remodeling of the synaptic nanoorganization. Two-color STED microscopy opens novel avenues for the *in vivo* investigation of the synaptic nanoorganization and its dynamics in the living brain. In the future this approach can be extended to study the plasticity of nanocolumns after experience, as well as over longer time-intervals in chronic experiments (Steffens et al., 2021) and different brain regions..

## MATERIALS AND METHODS

### DNA constructs

The transcriptionally regulated antibody-like protein to label PSD95 was obtained via plasmid pAAV-ZFN-hSyn-DIO-PSD95-FingR-Citrine-CCR5TC, with a double-floxed inverted open reading frame (DIO) as described in (Willig et al., 2021).

The plasmid pAAV-hSyn-DIO-myr-EGFP-LDLR(Ct) for dendritic membrane labelling was cloned as follows: First, we PCR amplified a myristoylation (myr) site-attached EGFP including the myristoylation sequence ATGGGCTGTGTGCAATGTAAGGATAAAGAAGCAACAAAACTGACG in the forward primer. Second, we split the C-terminal (Ct) cytoplasmic domains of low density lipoprotein receptor (LDLR; GenBank: AF425607, amino acid residues 813–862) (Kameda et al., 2008), finally designated as LDLR(Ct), into two parts (part 1 and part 2), designed 5’ phosphorylated forward and reverse primers for each part and hybridized each pair (Table 1). In the third and final step, the endonuclease-digested myr-EGFP PCR together with the LDLR(Ct) part1 and LDLR(Ct) part 2 were ligated into plasmid pAAV-hSyn-DIO-EYFP digested with AscI and NheI to finally obtain pAAV-hSyn-DIO-myrEGFP-LDLR(Ct).

**Table 1:**
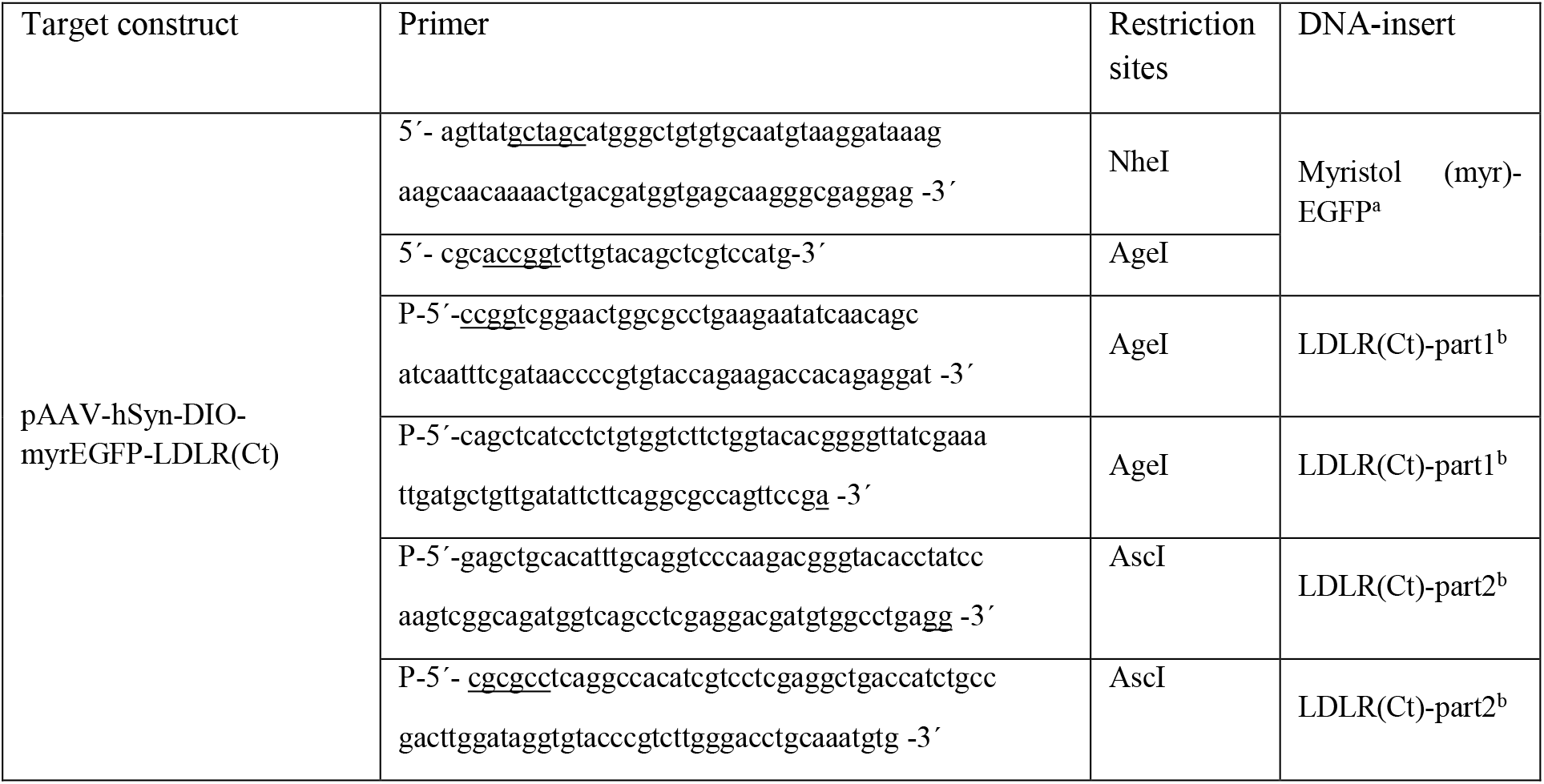
Overview of the primers and endonucleases. a: produced by PCR, b: generated by hybridization, P: phosphorylated; underlined nucleotides: restriction sites or part of them

Generation of the Cre recombinase expression plasmid pAAV-hSyn-Cre was performed as described (Wegner et al., 2017).

The crosstalk was determined by expression of a fusion protein consisting of the small peptide Lifeact (LA), that directly binds to F-actin, and a fluorescent protein: pAAV-hSyn-LA-EYFP and pAAV-hSyn-LA-EGFP (Willig et al., 2021, 2014).

### Virus production

Recombinant AAV particles with mixed serotype 1 and 2 of the pAAV plasmids encoding the proteins of interest were produced in HEK293-FT cells. The entire procedure is described in detail in (Wegner et al., 2017) and is applied here with the following modifications: After DNaseI treatment (30 min at 37 °C), the suspension was centrifuged at 1200 g for 10 min. The supernatant was filtered through an 0.45 μm sterile filter (Merck/Millipore, Darmstadt, Germany) and first applied to an Amicon Ultra-15, MWCO 100 kDa, centrifugal filter unit (Merck/Millipore, Darmstadt, Germany), followed by a Vivaspin^®^ 500, MWCO 100 kDa, centrifugal concentrator (Sartorius, Göttingen, Germany) to wash the virus in Opti-MEM™ Medium (ThermoFisher Scientific, Darmstadt, Germany) and concentrate to a final volume of 150 μl.

### Animals

All animal experiments were performed with C57BL/6J female mice reared at the animal facility of the Max Planck Institute of Experimental Medicine in Göttingen and housed with a 12-h light/dark cycle, with food and water available *ad libitum*. Experiments were performed according to the guidelines of the national law regarding animal protection procedures and were approved by the responsible authorities, the Niedersächsisches Landesamt für Verbraucherschutz (LAVES, identification number 33.9-42502-04-14/1463). All efforts were made to avoid animal suffering and to minimize the number of animals used.

### Housing conditions

Environmentally enriched (EE) animals were born and raised in the commercially available Marlau™ cage (Marlau™, Viewpoint, Lyon, France) (Fares et al., 2012), 580 x 400 x 320 mm in size including 2 floors, providing an extensive exploration area (Fig. S2). Three pregnant female mice were placed in an EE cage about 1 week before delivery. Pups were weaned and split by sex at post-natal day 30. The ground floor of the EE cage consisted of two separate compartments: the smaller part contained food and the larger part contained water, running wheels and a red house. To get food, mice had to use a stairway to reach the second floor where a maze was placed. After passing the maze, mice slided through a tunnel back to the ground floor, directly into the smaller compartment and had free access to food. By passing a one-way door they could enter the larger part of the ground floor to get water. To increase novelty and maintaining cognitive stimulation, the maze was changed 3 times per week with a total of 12 different configurations. Control mice (Ctr) were born and raised in standard cages of 365 x 207 x 140 mm in size. They were kept in 2–3 animals per cage, which were solely equipped with nesting material.

### AAV transduction

Adult (>12 weeks) female C57BL/6J mice were stereotaxically transduced with a mixture of AAV with mixed serotype 1 and 2. AAV1/2-ZFN-hSyn-DIO-PSD95-FingR-Citrine-CCR5TC, AAV1/2-hSyn-Cre and AAV1/2-hSyn-DIO-myrEGFP-LDLR(Ct) were stereotaxically injected at the same time into the visual cortex of the left hemisphere at a depth of ~500 μm below the pia transducing mainly layer 5 pyramidal neurons by using the following coordinates: 3.4 mm posterior to the bregma, 2.2 mm lateral to the midline, at an angle of 70° to the vertical, and with a feed forward of ~750 μm. Anesthesia was initiated by intraperitoneal injection of 0.05 mg/kg fentanyl, 5 mg/kg midazolam, and 0.5 mg/kg medetomidin (MMF). The mouse was head fixed with a stereotaxic frame and placed on a heating pad throughout the whole procedure to maintain a constant body temperature. The depth of the anesthesia was controlled by monitoring the pulse rate and O2 saturation with a pulse oximeter at the thigh of the mouse and body temperature with a rectal temperature probe. A gas mixture with a high O2 (47.5 vol%) and CO2 (2.5 vol%) content was applied to the mouse’s nose, significantly increasing the oxygenation. The eyes of the mouse were covered with eye ointment and the head was disinfected with 70% ethanol. The skin was cut by a ~ 0.5 cm long incision and a drop of local anesthetic (0.2 mg mepivacaine) was applied. Then, a 0.5 mm hole was drilled into the skull. 150 nl of AAV containing solution was pressure injected through a glass micropipette attached to a microinjector (Picospritzer^®^ III, Parker Hannifin Corp, Cleveland, Ohio) with an injection rate of ~50 nl per minute. After injection, the pipette was kept at the target location for an additional 2 minutes to allow the virus to disperse away. After retracting the micropipette, the incision was closed with a suture. Anesthesia was then antagonized by intraperitoneal administration of 0.1 mg/kg buprenorphine and 2.5 mg/kg atipamezole. The mouse was kept in a separate cage until full recovery and then put back into its original cage, and group-housed in the animal facility until the final experiment.

### Craniotomy for *in vivo* STED microscopy

Three to six weeks after viral transduction, a craniotomy was performed as described previously (Steffens et al., 2020). In brief, the mouse was anaesthetized with MMF, placed on a heating pad and vital functions and depth of anesthesia were controlled throughout the final experiment as described above. The head was mounted in a stereotaxic frame. The scalp was removed and a flat head bar was glued to the right hemisphere to leave enough space for the cranial window above the visual cortex, rostral to the former viral injection site. After drilling a circular groove (~2–3 mm) into the skull, the bony plate was carefully removed without causing a trauma. The remaining dura and arachnoid mater were carefully removed with a fine forceps. A small tube was positioned at the edge of the opening to drain excess cerebrospinal fluid if necessary. The craniotomy was sealed with a 6 mm diameter cover glass glued to the skull. The mouse was mounted on a tiltable plate which can be aligned perpendicular to the optical axis of the microscope in a quick and easy routine before being placed under the microscope (Steffens et al., 2020).

### Culture of primary hippocampal neurons, transduction and imaging

Primary cultures of rat hippocampal neurons were prepared from P0-P1 Wistar rats (RjHan:WI; The Jackson Laboratory, Bar Harbor, ME) of both sexes according to (D’Este et al., 2017). Neurons were cultured at 37 °C in a humidified atmosphere with 5% CO2 and transduced between 8 and 10 days *in vitro* (DIV) with the respective AAVs. After an incubation time of 7 days, live cell STED imaging was performed at room temperature.

### *In vivo* two-color STED microscope

A previously described custom-designed STED microscope (Wegner et al., 2018; Willig et al., 2014) was modified to accommodate for virtually crosstalk free two-color imaging as follows (Fig. 1). Excitation light (Exc1) provided by a pulsed laser diode emitting blue light at 483 nm (PiLas, Advanced Laser Diode Systems, Berlin, Germany) was complemented by a second excitation beam (Exc2). The beam of a Ti:Sapphire laser (MaiTai; Spectra-Physics, Santa Clara, CA) was split into two, pumping an optical parametric oscillator (OPO; APE, Berlin, Germany) emitting 80 MHz pulses at 595 nm for stimulated emission depletion (STED) and a supercontinuum device (FemtoWHITE800, NKT photonics, Birkerød, Denmark) generating white light. The white light was spectrally filtered for green light with a bandpass filter (BP1, BrightLine HC 520/5, Semrock, IDEX Health & Science, Rochester, NY) for selective excitation of EYFP or Citrine at 520 nm (Exc2, Fig. 1A,B). Exc1, Exc2 and the STED beam were co-aligned with dichroic mirrors. After passing a Yanus scan head (Till Photonics-FEI, Gräfelfing, Germany) consisting of two galvanometric mirrors for x-/y-scanning and relay optics, the three beams were focused by a 1.3 numerical aperture objective lens (PL APO, 63x, glycerol; Leica, Wetzlar, Germany). Additionally, the STED beam was passing a vortex phase plate (VPP; RPC Photonics, Rochester, NY) to create a doughnut shaped focal intensity pattern featuring zero intensity in the center. Temporal overlap of all three pulsed laser beams was achieved electronically by synchronizing the blue laser diode, Exc1, to the MaiTai and optically by an optical delay line for Exc2. Z-scanning was performed by moving the objective with a piezo (MIPOS 100PL; piezosystem jena GmbH, Jena, Germany). The back-projected fluorescence light was split at 515 nm with a dichroic mirror (Dm2, ZT502rdc-UF3; Chroma Technology Corporation, Bellow Falls, VT) into two beams. The shorter wavelength was reflected, filtered by a bandpass filter (BP3, BrightLine HC 504/12; Semrock) and focused onto a multimode fibre for confocal detection connected to an avalanche photodiode (APD, Excelitas, Waltham, MA). The transmitted, longer wavelength fluorescence light was also filtered with a bandpass filter (BP2, H544/23, AHF analysentechnik, Tübingen, Germany) and detected with an APD, respectively.

### In vivo two-color STED imaging of anesthetized mice

The *in vivo* STED microscopy was performed in layer 1 of the visual cortex at a depth of 5−20 μm below the pia. Spherical aberrations due to the tissue penetration were corrected on the first order by adapting the correction collar of the glycerol immersion objective for the best image quality at each field of view. Both excitation colors were alternated line-by-line; i.e. a line was recorded by excitation with blue light (Exc1) and then the same line was recorded again with green excitation light (Exc2). Potential drift between images and movement of the spine or PSD95 was therefore negligible. For imaging we picked different fields of view (FOV) with dendrites parallel to the focal plane. After recording a two-color STED image stack, the position of the motorized micrometer stage (MS-2000, Applied scientific instruments, Eugene, OR) was noted and an overview image taken. This process was repeated several times for different positions. After 30 min the micrometer stage was moved back to the first position. The position was confirmed by recording coarse overview images of the dendrite and thereby adjusting the z-position to the right depth. STED images were recorded as z-stacks of ~20–40 μm in x and y with 500 nm axial steps at different positions of the cranial window. All positions were repeatedly imaged 3–4 times at intervals of 30 min to 2 hours. Therefore, some dendritic regions were investigated at the time points 0 min, 30 min and 60 min, while other dendritic region within the cranial window were examined at the time points 0 min, 60 min and 120 min or even at 0 min, 120 min and 240 min. The benefit of a membrane label over the often used volume label is that the brightness of the spines and dendrites is similar whereas with a volume label spine heads and dendrites often outshine the small spine neck; thus, either the much darker neck is not visible between the bright head and dendrite or the detector is saturated at the bright heads and dendrites.

All images were recorded with a pixel dwell time of 5 μs, a pixel size of 30 x 30 nm in x and y, and z-stacks of 500 nm step size. Blue and green excitation power was 4.5 μW, respectively, in the back aperture of the objective. The average STED power in the back aperture was 37–45 mW for static images and 15 mW for time-lapse imaging.

### Data processing and analysis

Confocal and STED images were acquired by the software Imspector (Abberior Instruments, Göttingen, Germany). Analysis of size, shape, and brightness was performed manually in Fiji (Schindelin et al., 2012). For spine head and PSD95 analysis, the first and last plane were omitted to ensure that the spine (or PSD95, respectively) was located completely in the focal plane. We only analyzed spines that also bore a PSD and we only analyzed spines that extended from the dendrite mainly parallel to the focal plane. PSDs directly on the dendrite most likely representing shaft synapses and spines pointing upwards or downwards that could not be clearly resolved were not analyzed. First, images of each channel were processed as follows (Fiji commands shown in capital letters): 1. Smoothing: PROCESS > SMOOTH twice. 2. Brightness adjustment: IMAGE > ADJUST > BRIGHTNESS/CONTRAST set minimum to 1 instead of zero. 3. Overlay both one-color images: IMAGE > COLOR > MERGE CHANNELS. 4. Open ROI manager: ANALYSE > TOOLS > ROI MANAGER. Spine heads and PSD95 assemblies were encircled at their largest extent as shown in Fig. 2A to compute their area. Perforations or cluster of PSD95 were encircled to include only PSD95. For the analysis of brightness, PSD95 images were processed as described above. Using the ELLIPTICAL SELECTION tool, each PSD95 assembly was encircled and the brightness was displayed by the variable "RawIntDen", which is the sum of the intensity values of all pixels in the selected area. Spine density was obtained by dividing the total number of spines minus one by the length of the parent dendrite between the first and last spine. The length was measured with the FREEHAND LINE tool, in Z PROJECTION (maximum intensity) images. Branched spines were counted once.

Absolute changes of spine head area or PSD95 assembly area between two time points, t and t+1, were calculated by ΔA_abs_=A(t+1)-A(t). ‘A’ denotes the spine head area or the PSD95 area, respectively. Normalized changes of spine head area or PSD95 area were computed by ΔA_norm_=(A(t+1)-A(t))/(A(t+1)+A(t)) and percentage changes by ΔA%=(A(t+1)-A(t))/A(t)*100%.

The principal component analysis (PCA) was performed with the built in MATLAB^®^ (MathWorks^®^, Natick, MA) function ‘princomp’.

Pearson’s correlation coefficient and linear regression were computed in GraphPad Prism (version 7.04, Graphpad Software, San Diego, CA). The cross-correlation function was computed for different time intervals Δt by

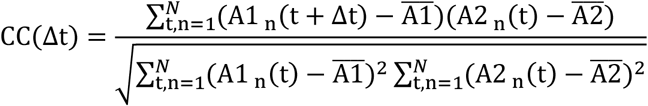

A1 and A2 denote spine head and PSD95 assembly size, respectively. N stands for the total number of spines, Δt the lag time and 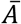 the average size of all spine heads, respectively PSD95 assemblies.

### Nanoorganization and morphological changes of PSD95

All non-macular PSD95 nanoorganizations were selected. Their shape was categorized by 3 blinded scientists into perforated or clustered shapes. Perforated PSDs were of continuous shape but complex such as a ring or horseshoe-like or more twisted. Clustered nanoorganizations were characterized by two or more separated assemblies of PSD95 per spine and the number of clusters was counted. Temporal changes in PSD95 morphology were analyzed as follows. Assemblies which did not alter their overall morphology or number of clusters were marked as ‘no change’. Assemblies showing minor modifications, such as a small movement of a sub-cluster, were classified as ‘subtle change’. Changes of the overall morphology such as a smooth continuous PSD falling apart into different clusters were characterized as ‘strong change’. All changes were referred to time point t = 0 min.

### Statistical analysis

Statistical analysis was performed in GraphPad Prism. Positively skewed data sets were compared by the Kolmogorov Smirnov test. Log-transformed data and normalized relative changes were normally distributed and compared by an unpaired *t*-test with Welch’s correction. The number of analyzed mice and spines, p-value, and the specific statistical test performed for each experiment are included in the appropriate figure legend. All tests were applied two-sided where applicable. Probabilities are symbolized by asterisks: *p < 0.05; **p < 0.01; ***p < 0.001, ****p < 0.0001

### Materials availability

This study did not generate new unique reagents.

### Data and materials availability

Image data files are provided by reasonable request through the corresponding author.

## Acknowledgments

We thank Dr. S. Löwel (Göttingen University) for suggesting and providing the Marlau™ cages, the animal facility of the MPI for Experimental Medicine for excellent support and Dr. Karl Deisseroth and Dr. Don Arnold for providing plasmids. This work was funded by the Deutsche Forschungsgemeinschaft (DFG, German Research Foundation) within the DFG Research Center and Cluster of Excellence (EXC 171, Area A1) “Nanoscale Microscopy and Molecular Physiology of the Brain” (WW, HS, KIW) and under Germany’s Excellence Strategy - EXC 2067/1-390729940 (KIW, CG).

## Author contributions

Conceptualization (WW, KIW); STED microscope design and construction (KIW); Cloning and virus production (WW, CG); Surgery (HS, WW); Imaging (WW, KIW); Analysis (WW, KIW, FW); Writing (WW, KIW).

## Conflict of interest

The authors declare no competing interests.

## Notes

### Competing Interest Statement

The authors have declared no competing interest.

## References

Arellano JI, Benavides-Piccione R, Defelipe J, Yuste R. 2007. Ultrastructure of dendritic spines: correlation between synaptic and spine morphologies. Front Neurosci 1:131–43. doi:10.3389/neuro.01.1.1.010.2007

Artola A, von Frijtag JC, Fermont PCJ, Gispen WH, Schrama LH, Kamal A, Spruijt BM. 2006. Long-lasting modulation of the induction of LTD and LTP in rat hippocampal CA1 by behavioural stress and environmental enrichment. Eur J Neurosci 23:261–272. doi:10.1111/j.1460-9568.2005.04552.x

Baroncelli L, Sale A, Viegi A, Maya Vetencourt JF, De Pasquale R, Baldini S, Maffei L. 2010. Experience-dependent reactivation of ocular dominance plasticity in the adult visual cortex. Exp Neurol 226:100–109. doi:10.1016/j.expneurol.2010.08.009

Bosch M, Castro J, Saneyoshi T, Matsuno H, Sur M, Hayashi Y. 2014. Structural and molecular remodeling of dendritic spine substructures during long-term potentiation. Neuron 82:444–459. doi:10.1016/j.neuron.2014.03.021

Bottanelli F, Kromann EB, Allgeyer ES, Erdmann RS, Baguley SW, Sirinakis G, Schepartz A, Baddeley D, Toomre DK, Rothman JE, Bewersdorf J. 2016. Two-colour live-cell nanoscale imaging of intracellular targets. Nat Commun 7:1–5. doi:10.1038/ncomms10778

Buschler A, Manahan-Vaughan D. 2012. Brief environmental enrichment elicits metaplasticity of hippocampal synaptic potentiation in vivo. Front Behav Neurosci 6:1–10. doi:10.3389/fnbeh.2012.00085

Cane M, Maco B, Knott G, Holtmaat A. 2014. The Relationship between PSD-95 Clustering and Spine Stability In Vivo. J Neurosci 34:2075–2086. doi:10.1523/JNEUROSCI.3353-13.2014

Chazeau A, Giannone G. 2016. Organization and dynamics of the actin cytoskeleton during dendritic spine morphological remodeling. Cell Mol Life Sci 73:3053–3073. doi:10.1007/s00018-016-2214-1

Chen H, Tang AH, Blanpied TA. 2018. Subsynaptic spatial organization as a regulator of synaptic strength and plasticity. Curr Opin Neurobiol 51:147–153. doi:10.1016/j.conb.2018.05.004

Compans B, Choquet D, Hosy E. 2016. Review on the role of AMPA receptor nano-organization and dynamic in the properties of synaptic transmission. Neurophotonics 3:041811. doi:10.1117/1.NPh.3.4.041811

Cook SG, Goodell DJ, Restrepo S, Arnold DB, Bayer KU. 2019. Simultaneous Live Imaging of Multiple Endogenous Proteins Reveals a Mechanism for Alzheimer’s-Related Plasticity Impairment. Cell Rep 27:658–665.e4. doi:10.1016/j.celrep.2019.03.041

D’Este E, Kamin D, Balzarotti F, Hell SW. 2017. Ultrastructural anatomy of nodes of Ranvier in the peripheral nervous system as revealed by STED microscopy. Proc Natl Acad Sci 114:E191–E199. doi:10.1073/pnas.1619553114

Eggeling C, Willig KI, Sahl SJ, Hell SW. 2015. Lens-based fluorescence nanoscopy. Q Rev Biophys 48:178–243. doi:10.1017/s0033583514000146

El-Husseini A, Schnell E, Chetkovich D. 2000. PSD-95 involvement in maturation of excitatory synapses. Science 290:1364–8. doi:10.1126/science.290.5495.1364

Fares RP, Kouchi H, Bezin L. 2012. Standardized environmental enrichment for rodents in Marlau cage. Protoc Exch 1–14. doi:10.1038/protex.2012.036

Franks KM, Stevens CF, Sejnowski TJ. 2003. Independent Sources of Quantal Variability at Single Glutamatergic Synapses. J Neurosci 23:3186–3195. doi:10.1523/JNEUROSCI.23-08-03186.2003

Gelfo F, De Bartolo P, Giovine A, Petrosini L, Leggio MG. 2009. Layer and regional effects of environmental enrichment on the pyramidal neuron morphology of the rat. Neurobiol Learn Mem 91:353–365. doi:10.1016/j.nlm.2009.01.010

Göttfert F, Wurm CA, Mueller V, Berning S, Cordes VC, Honigmann A, Hell SW. 2013. Coaligned dual-channel STED nanoscopy and molecular diffusion analysis at 20 nm resolution. Biophys J 105:6–8. doi:10.1016/j.bpj.2013.05.029

Gross GG, Junge JA, Mora RJ, Kwon H-BB, Olson CA, Takahashi TT, Liman ER, Ellis-Davies GCRR, McGee AW, Sabatini BL, Roberts RW, Arnold DB. 2013. Recombinant Probes for Visualizing Endogenous Synaptic Proteins in Living Neurons. Neuron 78:971–985. doi:10.1016/j.neuron.2013.04.017

Harris K, Jensen F, Tsao B. 1992. Three-dimensional structure of dendritic spines and synapses in rat hippocampus (CA1) at postnatal day 15 and adult ages: implications for the maturation of synaptic physiology and long-term potentiation [published erratum appears in J Neurosci 1992 Aug;1. J Neurosci 12:2685–2705. doi:10.1523/JNEUROSCI.12-07-02685.1992

Harris KM, Weinberg RJ. 2012. Ultrastructure of Synapses in the Mammalian Brain. Cold Spring Harb Perspect Biol 4:a005587–a005587. doi:10.1101/cshperspect.a005587

Hazan L, Ziv NE. 2020. Activity Dependent and Independent Determinants of Synaptic Size Diversity. J Neurosci 40:2828–2848. doi:10.1523/JNEUROSCI.2181-19.2020

Herring BE, Nicoll RA. 2016. Long-Term Potentiation: From CaMKII to AMPA Receptor Trafficking. Annu Rev Physiol 78:351–365. doi:10.1146/annurev-physiol-021014-071753

Hofer SB, Mrsic-Flogel TD, Bonhoeffer T, Hübener M. 2009. Experience leaves a lasting structural trace in cortical circuits. Nature 457:313–7. doi:10.1038/nature07487

Holler S, Köstinger G, Martin KAC, Schuhknecht GFP, Stratford KJ. 2021. Structure and function of a neocortical synapse. Nature. doi:10.1038/s41586-020-03134-2

Hruska M, Henderson N, Le Marchand SJ, Jafri H, Dalva MB. 2018. Synaptic nanomodules underlie the organization and plasticity of spine synapses. Nat Neurosci 21:1. doi:10.1038/s41593-018-0138-9

Kalogeraki E, Pielecka-Fortuna J, Löwel S. 2017. Environmental enrichment accelerates ocular dominance plasticity in mouse visual cortex whereas transfer to standard cages resulted in a rapid loss of increased plasticity. PLoS One 12:1–24. doi:10.1371/journal.pone.0186999

Kameda H, Furuta T, Matsuda W, Ohira K, Nakamura K, Hioki H, Kaneko T. 2008. Targeting green fluorescent protein to dendritic membrane in central neurons. Neurosci Res 61:79–91. doi:10.1016/j.neures.2008.01.014

Lee HK, Kirkwood A. 2019. Mechanisms of Homeostatic Synaptic Plasticity in vivo. Front Cell Neurosci 13:1–7. doi:10.3389/fncel.2019.00520

Leggio MG, Mandolesi L, Federico F, Spirito F, Ricci B, Gelfo F, Petrosini L. 2005. Environmental enrichment promotes improved spatial abilities and enhanced dendritic growth in the rat. Behav Brain Res 163:78–90. doi:10.1016/j.bbr.2005.04.009

Loewenstein Y, Kuras A, Rumpel S. 2011. Multiplicative dynamics underlie the emergence of the log-normal distribution of spine sizes in the neocortex in vivo. J Neurosci 31:9481–8. doi:10.1523/JNEUROSCI.6130-10.2011

MacGillavry HD, Song Y, Raghavachari S, Blanpied TA. 2013. Nanoscale scaffolding domains within the postsynaptic density concentrate synaptic ampa receptors. Neuron 78:615–622. doi:10.1016/j.neuron.2013.03.009

Mandolesi L, Gelfo F, Serra L, Montuori S, Polverino A, Curcio G, Sorrentino G. 2017. Environmental Factors Promoting Neural Plasticity: Insights from Animal and Human Studies. Neural Plast 2017:7219461. doi:https://dx.doi.org/10.1155/2017/7219461

Matsuzaki M, Honkura N, Ellis-Davies GCR, Kasai H. 2004. Structural basis of long-term potentiation in single dendritic spines. Nature 429:761–766. doi:10.1038/nature02617

Meyer D, Bonhoeffer T, Scheuss V. 2014. Balance and Stability of Synaptic Structures during Synaptic Plasticity. Neuron 82:430–443. doi:10.1016/j.neuron.2014.02.031

Moneron G, Hell SW. 2009. Two-photon excitation STED microscopy. Opt Express 17:14567. doi:10.1364/oe.17.014567

Mongillo G, Rumpel S, Loewenstein Y. 2017. Intrinsic volatility of synaptic connections — a challenge to the synaptic trace theory of memory. Curr Opin Neurobiol 46:7–13. doi:10.1016/j.conb.2017.06.006

Nagerl U V., Willig KI, Hein B, Hell SW, Bonhoeffer T. 2008. Live-cell imaging of dendritic spines by STED microscopy. Proc Natl Acad Sci 105:18982–18987. doi:10.1073/pnas.0810028105

Nithianantharajah J, Hannan AJ. 2006. Enriched environments, experience-dependent plasticity and disorders of the nervous system. Nat Rev Neurosci 7:697–709. doi:10.1038/nrn1970

Nithianantharajah J, Levis H, Murphy M. 2004. Environmental enrichment results in cortical and subcortical changes in levels of synaptophysin and PSD-95 proteins. Neurobiol Learn Mem 81:200–10. doi:10.1016/j.nlm.2004.02.002

Panatier A, Arizono M, Nagerl U V. 2014. Dissecting tripartite synapses with STED microscopy. Philos Trans R Soc B Biol Sci 369:20130597–20130597. doi:10.1098/rstb.2013.0597

Savtchenko LP, Rusakov DA. 2014. Moderate AMPA receptor clustering on the nanoscale can efficiently potentiate synaptic current. Philos Trans R Soc B Biol Sci 369. doi:10.1098/rstb.2013.0167

Schindelin J, Arganda-Carreras I, Frise E, Kaynig V, Longair M, Pietzsch T, Preibisch S, Rueden C, Saalfeld S, Schmid B, Tinevez J-Y, White DJ, Hartenstein V, Eliceiri K, Tomancak P, Cardona A. 2012. Fiji: an open-source platform for biological-image analysis. Nat Methods 9:676–682. doi:10.1038/nmeth.2019

Sigler A, Oh WC, Imig C, Altas B, Kawabe H, Cooper BH, Kwon HB, Rhee JS, Brose N. 2017. Formation and Maintenance of Functional Spines in the Absence of Presynaptic Glutamate Release. Neuron 94:304–311.e4. doi:10.1016/j.neuron.2017.03.029

Statman A, Kaufman M, Minerbi A, Ziv NE, Brenner N. 2014. Synaptic Size Dynamics as an Effectively Stochastic Process. PLoS Comput Biol 10. doi:10.1371/journal.pcbi.1003846

Steffens H, Mott AC, Li S, Wegner W, Švehla P, Kan VWY, Wolf F, Liebscher S, Willig KI. 2021. Stable but not rigid: Chronic in vivo STED nanoscopy reveals extensive remodeling of spines, indicating multiple drivers of plasticity. Sci Adv 7:eabf2806. doi:10.1126/sciadv.abf2806

Steffens H, Wegner W, Willig KI. 2020. In vivo STED microscopy: A roadmap to nanoscale imaging in the living mouse. Methods 174:42–48. doi:10.1016/j.ymeth.2019.05.020

Stein V, House DRC, Bredt DS, Nicoll RA. 2003. Postsynaptic Density-95 Mimics and Occludes Hippocampal Long-Term Potentiation and Enhances Long-Term Depression. J Neurosci 23:5503–5506. doi:10.1523/JNEUROSCI.23-13-05503.2003

Stewart MG, Medvedev NI, Popov VI, Schoepfer R, Davies H a., Murphy K, Dallérac GM, Kraev I V., Rodríguez JJ. 2005. Chemically induced long-term potentiation increases the number of perforated and complex postsynaptic densities but does not alter dendritic spine volume in CA1 of adult mouse hippocampal slices. Eur J Neurosci 21:3368–3378. doi:10.1111/j.1460-9568.2005.04174.x

Tang A-H, Chen H, Li TP, Metzbower SR, MacGillavry HD, Blanpied TA. 2016. A trans-synaptic nanocolumn aligns neurotransmitter release to receptors. Nature 536:210–4. doi:10.1038/nature19058

Tønnesen J, Katona G, Rózsa B, Nägerl UV. 2014. Spine neck plasticity regulates compartmentalization of synapses. Nat Neurosci 17:678–85. doi:10.1038/nn.3682

Tønnesen J, Nadrigny F, Willig KI, Wedlich-Söldner R, Nägerl UV. 2011. Two-Color STED Microscopy of Living Synapses Using A Single Laser-Beam Pair. Biophys J 101:2545–2552. doi:10.1016/j.bpj.2011.10.011

Villa KL, Berry KP, Subramanian J, Cha JW, Oh WC, Kwon HB, Kubota Y, So PTC, Nedivi E. 2016. Inhibitory Synapses Are Repeatedly Assembled and Removed at Persistent Sites In Vivo. Neuron 89:756–769. doi:10.1016/j.neuron.2016.01.010

Wegner W, Ilgen P, Gregor C, van Dort J, Mott AC, Steffens H, Willig KI. 2017. In vivo mouse and live cell STED microscopy of neuronal actin plasticity using far-red emitting fluorescent proteins. Sci Rep 7:11781. doi:10.1038/s41598-017-11827-4

Wegner W, Mott AC, Grant SGN, Steffens H, Willig KI. 2018. In vivo STED microscopy visualizes PSD95 sub-structures and morphological changes over several hours in the mouse visual cortex. Sci Rep 8:219. doi:10.1038/s41598-017-18640-z

Willig KI, Kellner RR, Medda R, Hein B, Jakobs S, Hell SW. 2006. Nanoscale resolution in GFP-based microscopy. Nat Methods 3:721–723. doi:10.1038/nmeth922

Willig KI, Steffens H, Gregor C, Herholt A, Rossner MJ, Hell SW. 2014. Nanoscopy of Filamentous Actin in Cortical Dendrites of a Living Mouse. Biophys J 106:L01–L03. doi:10.1016/j.bpj.2013.11.1119

Willig KI, Wegner W, Müller A, Clavet-Fournier V, Steffens H. 2021. Multi-label in vivo STED microscopy by parallelized switching of reversibly switchable fluorescent proteins. Cell Rep 35:109192. doi:10.1016/j.celrep.2021.109192

Yuste R, Bonhoeffer T. 2001. Morphological Changes in Dendritic Spines Associated with Long-Term Synaptic Plasticity. Annu Rev Neurosci 24:1071–1089. doi:10.1146/annurev.neuro.24.1.1071

Ziv NE, Brenner N. 2018. Synaptic Tenacity or Lack Thereof: Spontaneous Remodeling of Synapses. Trends Neurosci 41:89–99. doi:10.1016/j.tins.2017.12.003

Ziv NE, Fisher-Lavie A. 2014. Presynaptic and Postsynaptic Scaffolds. Neurosci 20:439–452. doi:10.1177/1073858414523321

