## Supplementary data for "Environmental enrichment enhances patterning and remodeling of synaptic nanoarchitecture revealed by STED nanoscopy"

### Supplementary Material

#### Supplementary Figures

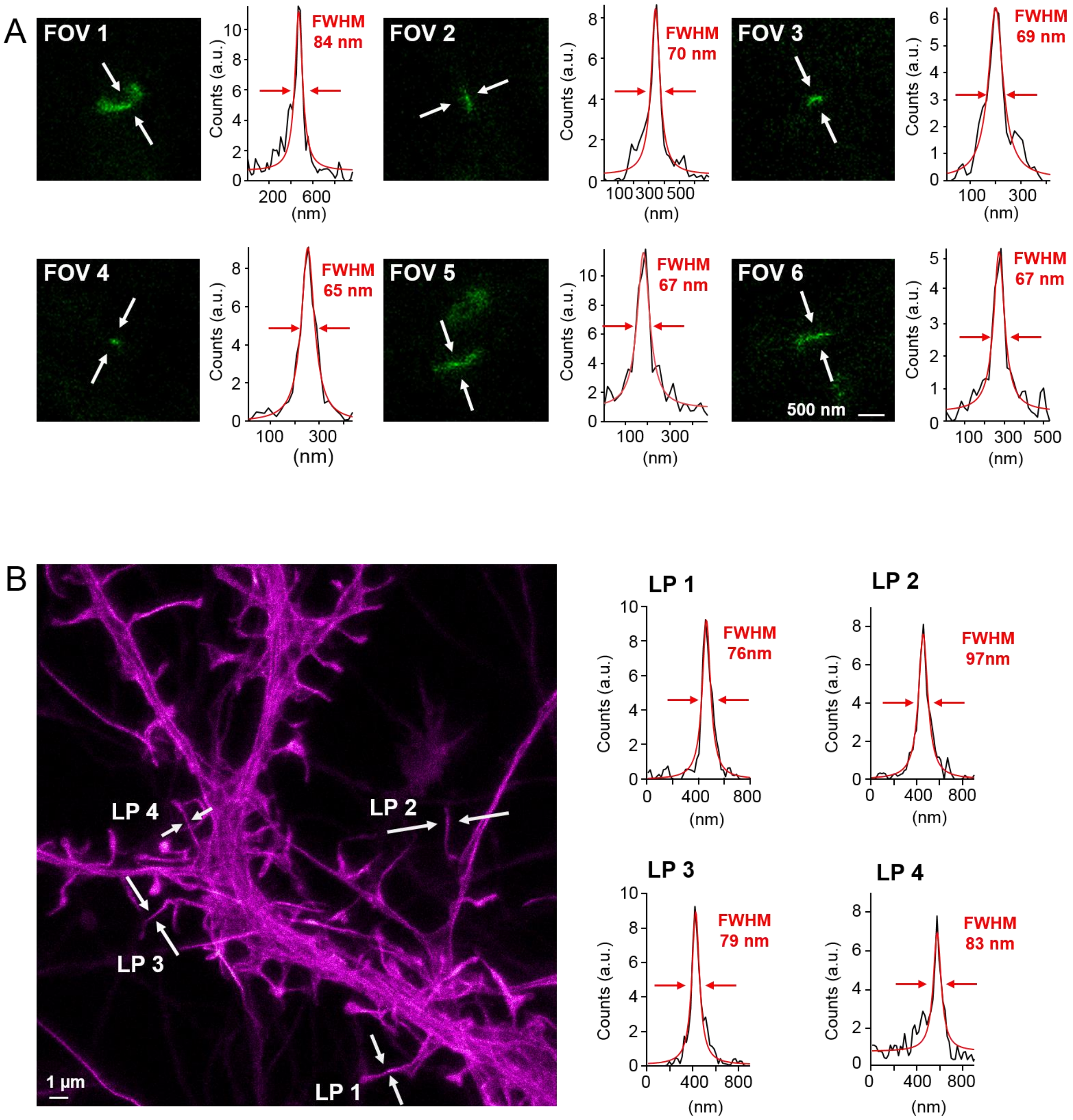


**Figure S1: Resolving capability of the custom-built two-color *in vivo* STED microscope. (A)** *In vivo* measurement of PSD95-FingR-Citrine. Fields of view (FOV) represent a selection of magnified PSD95 assemblies (green) from different STED image stacks. The resolving power was determined by fitting a Lorentzian function (red line) to the line profile (black line) which was averaged over 3 pixels at the indicated position (white arrows). The average of all determined full-widths at half-maximum (FWHM) is 70 nm, which is the upper estimate of the resolution of the custom-built STED microscope for Citrine. It is measured in the actual experiment and includes all potential distortions due to the tissue penetration of the excitation and STED beams and potential movement. **(B)** Live-cell STED microscopy of Lifeact-EGFP in neuronal cultures to determine the resolving power of the STED microscope for EGFP. 4 line profiles (LP) were taken at different positions as described in (A) which are in average 84 nm in FWHM.

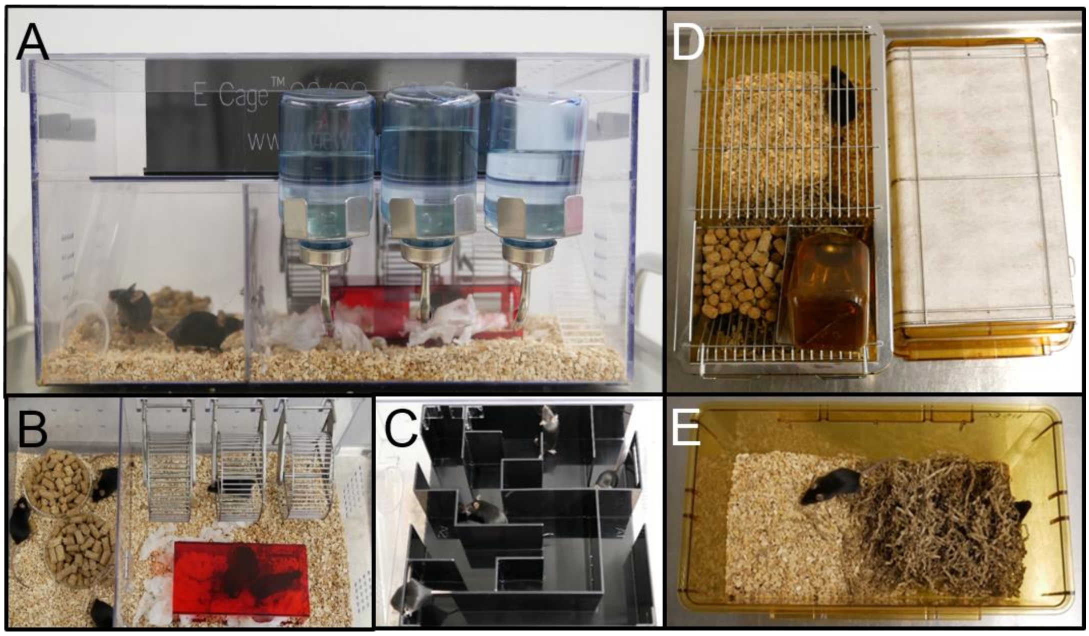


**Figure S2: Housing conditions for enriched environment (EE) and control (Ctr) mice. (A-C)** Standardized environmental enrichment cage design in the commercially available Marlau™ cage; dimension: 570 x 370 x 320 mm, 2 floors connected by a ladder and a tube. The cage contains 3 running-wheels, a red house, nesting material (B) and on the second floor a maze (C), which is changed 3 times a week. **(D, E)** Control mice are raised in a standard cage of 365 x 207 x 140 mm (floor area 530 cm$^2$) with nesting material only.

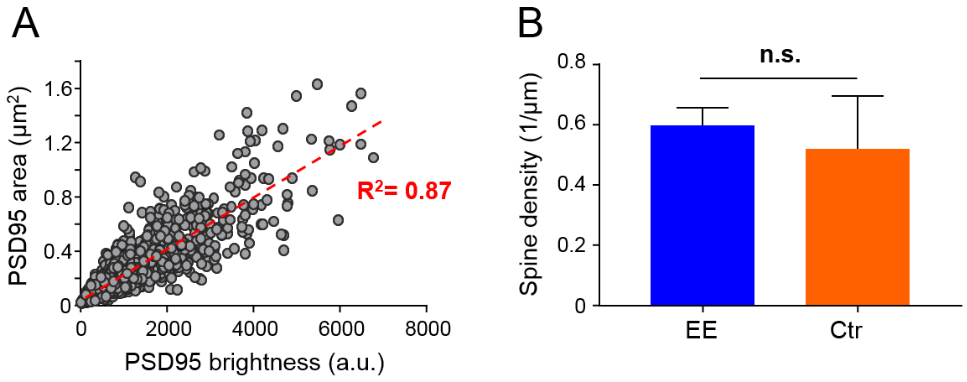


**Figure S3: (A) Analysis of PSD95 brightness.** Correlation of all EE and Ctr PSD95 areas in µm$^2$ and corresponding PSD95 brightness in *arbitrary units* (a.u.). The red dashed line represents a linear regression fit. Data of 8 mice and in total n = 1763 PSDs. **(B) Spine density.** Spine density in layer 1 of the visual cortex is not significantly different between EE and Ctr. Bars show median + 95% CI (Mann-Whitney U test, $p = 0.68$). 4x ♀mice per group and $n_{Ctr} = 59$, $n_{EE} = 99$ dendrites were analyzed.

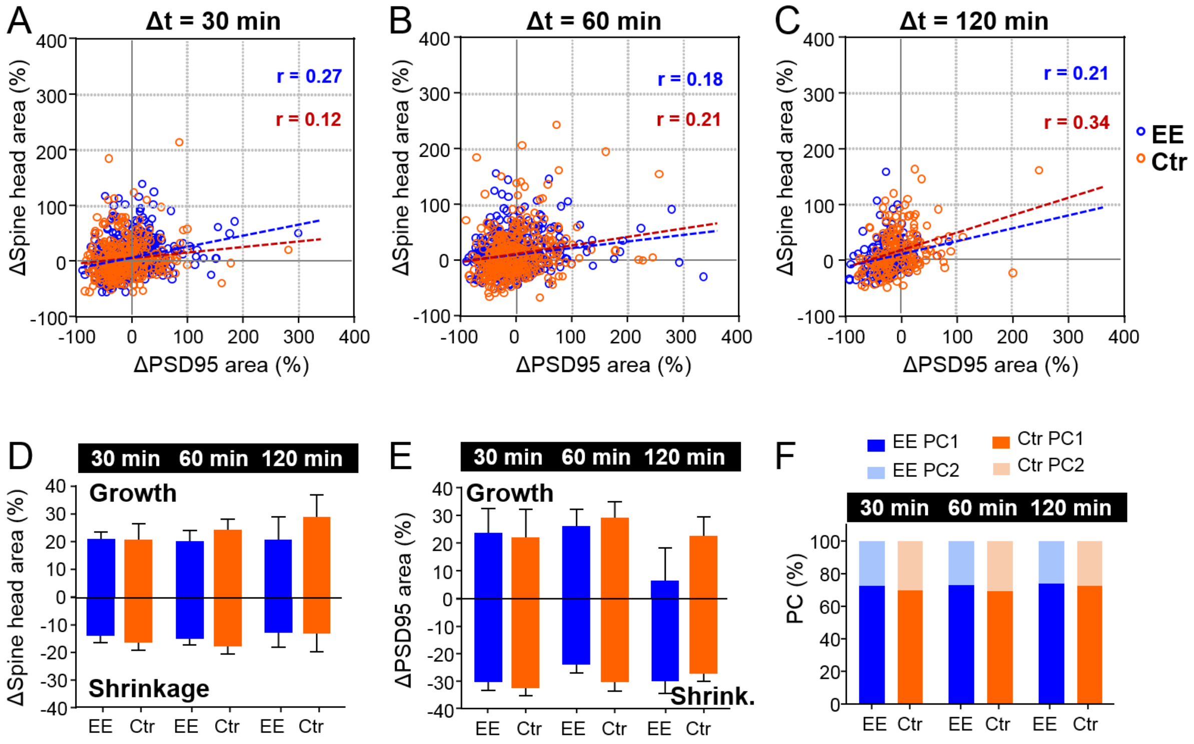


**Figure S4: Temporal changes in spine head and PSD95 area over time course**. **(A–C)** Scatter plot of percentage changes over time intervals of 30 min, 60 min and 120 min. Regression lines are dashed and Pearson's correlation coefficient *r* is displayed (deviation from zero: $p < 0.0001$). **(D, E)** Average percentage change of growing and shrinking spine head area (D) and PSD95 area (E). (D) Spine head growth: 30 min, EE: 21%, Ctr: 21%; 60 min, EE: 21%, Ctr: 24%; 120 min, EE: 21%, Ctr: 29% (Kruskal-Wallis with Dunn's multiple comparisons test; EE: 30 min vs. 60 min: $p > 0.99$ and vs. 120 min: $p > 0.99$, Ctr: 30 min vs. 60 min: $p > 0.99$ and vs 120 min: $p = 0.80$); Spine head shrinkage: 30 min, EE: -14%, Ctr: -17%; 60 min, EE: -15%, Ctr: -18%; 120 min, EE: -13%, Ctr: -13% (Kruskal-Wallis with Dunn's multiple comparisons test; EE: 30 min vs. 60 min: $p > 0.99$ and vs. 120 min: $p > 0.99$, Ctr: 30 min vs. 60 min: $p > 0.99$ and vs. 120 min: $p > 0.99$). Neither growth nor shrinkage shows a significant change over time. The average percentage spine head growth is not different between EE and Ctr (Kruskal-Wallis with Dunn's multiple comparisons test; spine head, EE vs. Ctr for 30 min, 60 min, 120 min: $p > 0.90$), neither is the average shrinkage (Kruskal-Wallis with Dunn's multiple comparisons test; spine head, EE vs. Ctr for 30 min, 60 min, 120 min: $p > 0.10$). (E) PSD95 growth: 30 min, EE: 24%, Ctr: 22%; 60 min, EE: 26%, Ctr: 29%; 120 min, EE: 6%, Ctr: 23%; PSD95 shrinkage: 30 min, EE: -30%, Ctr: -33%; 60 min, EE: -24%, Ctr: -30%; 120 min, EE: -30%, Ctr: -27%. Plots show median +95% CI. **(F)** Relative percentage of PC1 and PC2 of total variance. Data from Fig. 3H, I. (A–E) The same dataset as in Fig. 3D–F was analyzed.

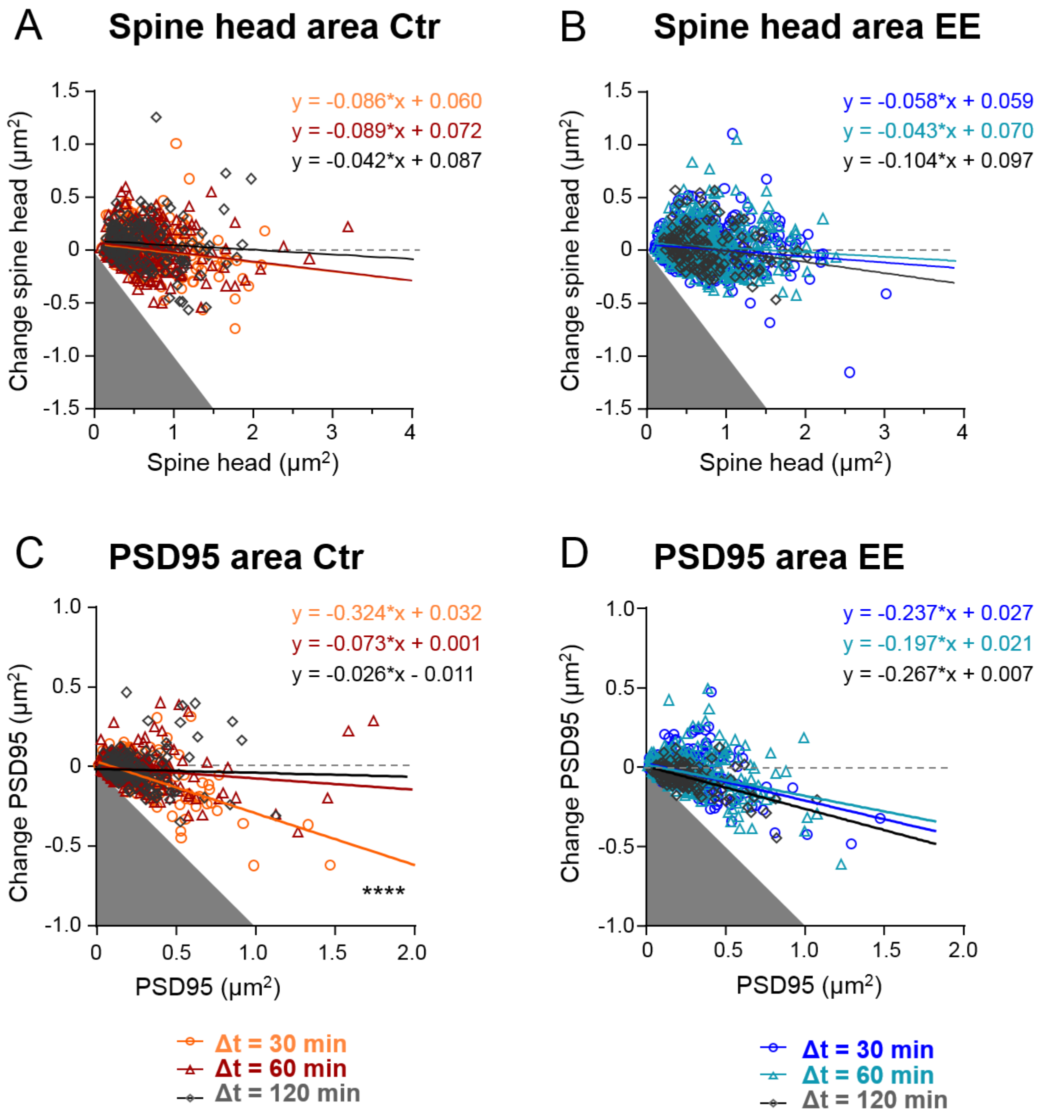


**Figure S5: Size changes of EE and Ctr housed mice. (A–D)** Changes in area after time intervals Δt of 30 min, 60 min and 120 min plotted as a function of their area at time point t for spine head area of Ctr housed mice (A), spine head area of EE housed mice (B), PSD95 area of Ctr housed mice (C) and PSD95 area of EE housed mice (D). Straight lines show linear regression fit of the displayed equation. Analysis of Covariance, Slopes equal? A: $p = 0.48$, B: $p = 0.35$, C: ****$p < 0.0001$, D: $p = 0.18$. Same data as Fig. 4.
